# STRUCT: a statistical approach to identify RNA secondary structures from raw sequencing data, bypassing multiple sequence alignment

**DOI:** 10.1101/2024.10.03.616574

**Authors:** Julie Fangran Wang, Arjun Rustagi, Julia Salzman

## Abstract

RNA secondary and tertiary structures are essential to life. Experimental methods to detect RNA structure, such as X-ray crystallography and chemical probing, are incisive but suffer from low throughput and dimensionality. Computational approaches, leveraging evolutionary signals from correlated mutations, provide an alternative means to infer RNA structures. However, these methods require assembly and face challenges due to statistical biases inherent in multiple sequence alignment (MSA). Furthermore, these methods cannot exploit a given RNA element’s full spectrum of natural sequence variations. Here, we introduce STRUCT (**S**tatistical **T**esting of **R**NA **U**nits with **C**ovariation **T**raits), an assembly-free, MSA-free, and metadata-free statistical method for identifying conserved RNA structures from raw sequencing data, quantifying base-pair covariations or stem variation exclusion in the putative RNA structures. We show STRUCT rediscovers known HIV structural elements and identifies conserved rRNA structures in metatranscriptomics samples. Moreover, STRUCT finds viral structures in mosquito metatranscriptomics samples *de novo*, including previously unannotated viral genomes, highlighting the method’s potential for viral discovery. STRUCT is an ultra-fast, easy-to-use, and robust tool that excels in high-throughput RNA structure prediction and hypothesis generation, presenting a novel approach for discovering structural RNA elements.

## 1 Backgroud

Highly conserved RNA secondary and tertiary structures are a hallmark of functional RNAs across life [1]. Examples include rRNA, tRNA, and the 7SL RNA and its derived SINE elements [2]. Many other examples, from ribozymes (e.g., the group I intron catalytic RNA) to riboswitches [3] to crRNA that direct Cas proteins to their targets [4] exhibit structural conservation. Experimentally, RNA structures can be determined directly (e.g., small angle X-ray scattering, X-ray crystallography, or cryogenic electron microscopy) or indirectly via biochemical probing (e.g., RNase footprinting [5], chemical mapping via SHAPE [6], and NGS methods [7]). These methods have proven to be very productive [8] but cannot scale with the present pace of biological sequence discovery [9]. While seminal developments have been made to identify and infer RNA secondary structures from sequence data [1, 10, 11], these methods still fail to keep pace with the sequence discovery rate.

Evolutionary signatures of biomolecular structures have proven to be potent signals for protein structure inference [12]. Analogously, comparative genomic analysis of orthologous RNA sequences can be used to infer RNA secondary structures. The first evidence for such an RNA structure signal came from the analysis of compensatory mutations in multiple sequence alignment [13]. The logic for this idea is as follows: assuming an RNA’s secondary structure is necessary for cellular fitness, a single base-pair-disrupting base change in such an RNA’s sequence will be deleterious to fitness; however, a compensating mutation will “rescue” the structure and have no or little net fitness cost. Thus, observing compensatory mutations (in the form of co-varying positions in a multiple sequence alignment of orthologous RNAs) in an area of inferred local structure can be both a signature of secondary structure and of selection for said structure. This idea has been explored extensively and adapted to experimental techniques for RNA secondary structure inference [1, 6, 14].

Rigorous analytic methods to infer RNA structure through multiple sequence alignment (MSA) have been developed and used to discover novel structural elements, most notably through tools such as R-scape [11]. These authors have also provided insights into the inferential pitfalls of some current methods based on MSA, including false positive calls of structural covariation [15]. First, MSA reduces the rich variation in microbial or even eukaryotic genome diversity, within and between hosts, to representative sequences. Some extensive compensatory mutations may be censored out of MSA. Second, these alignments are themselves computational procedures; thus, inference on them is conditional on the algorithmic output, that is, potentially statistically biased. This bias has been articulated in previous work [15] and includes (i) Misalignment or inclusion of non-homologous sequences can lead to incorrect conclusions; (ii) MSA inference may suffer from evolutionary confounding as described in a rich body of work: Phylogenetic relationships can introduce covariations that aren’t due to structural constraints. The existence of covariations as typically quantified must be considered conditional on evolutionary history [11]. Proposed methods seek to remove confounding by descent/evolutionary history. However, in the microbial world, phylogenetic trees cannot fully capture genetic diversity partly due to the prevalence of horizontal gene transfer [16], and to rapid mutation rates within hosts [17], in the case of RNA viruses. In summary, classical analysis of covariation is a conditional procedure biased by the process of genome assembly and MSA.

Critical obstacles inherent to existing multi-step bioinformatic processes preclude the high throughput and precise de novo computational prediction of RNA structures. Namely: 1) genomes or genome fragments must be assembled; 2) a multiple sequence alignment on them must be performed; and 3) a statistical decision and scoring of putative compensatory mutations must be performed without a closed form *p*-value and one that is confounded by phylogeny [11]. With rare exceptions (e.g., a genomic approach relying on a reference genome that yielded new riboswitches [18]), all of these steps have created a bottleneck for discovery. In addition, RNA structures within RNA viruses are critical with known functions (e.g., in poliovirus [19], SARS-CoV-2 [20], flavivirus [21]) that are relatively conserved in a background of rapid genomic variation. At the same time, RNA viruses are characterized by high mutation rates, exist as quasispecies [22], and can rapidly evolve within and between hosts, making assembly difficult. Raw sequences, not assemblies, are the evolutionary “tinder” of viruses and provide an insight into the “fitness landscape” they explore and which are selected.

Here, we introduce a direct statistical approach, STRUCT (https://github.com/juliefrwang/splash-structure), to detect conserved secondary structures from sequencing data(Figure 1A). The method quantifies compensatory mutations in stem-loops and detects cases of apparent negative selection against putatively deleterious structure-disrupting variants. STRUCT analyzes variable sequences linked to a shared fixed sequence from raw RNA-seq data, infers potential compensartory mutations under selection. By providing a direct statistical test for compensatory or absent mutaitons in stems, STRUCT detects RNA secondary structures bypassing MSA and its accompanying statistical and bioinformatic pitfalls[15]. STRUCT does not require a reference genome and enables statistical prioritization of sequences that exhibit higher levels of compensatory variation than a null distribution, providing efficiency and convenience over other RNA structure detection methods (Figure 1B). We show STRUCT provides precise *p*-values by theory and by simulation, as well as demonstrate its performance on real-world data.

**Fig. 1.**
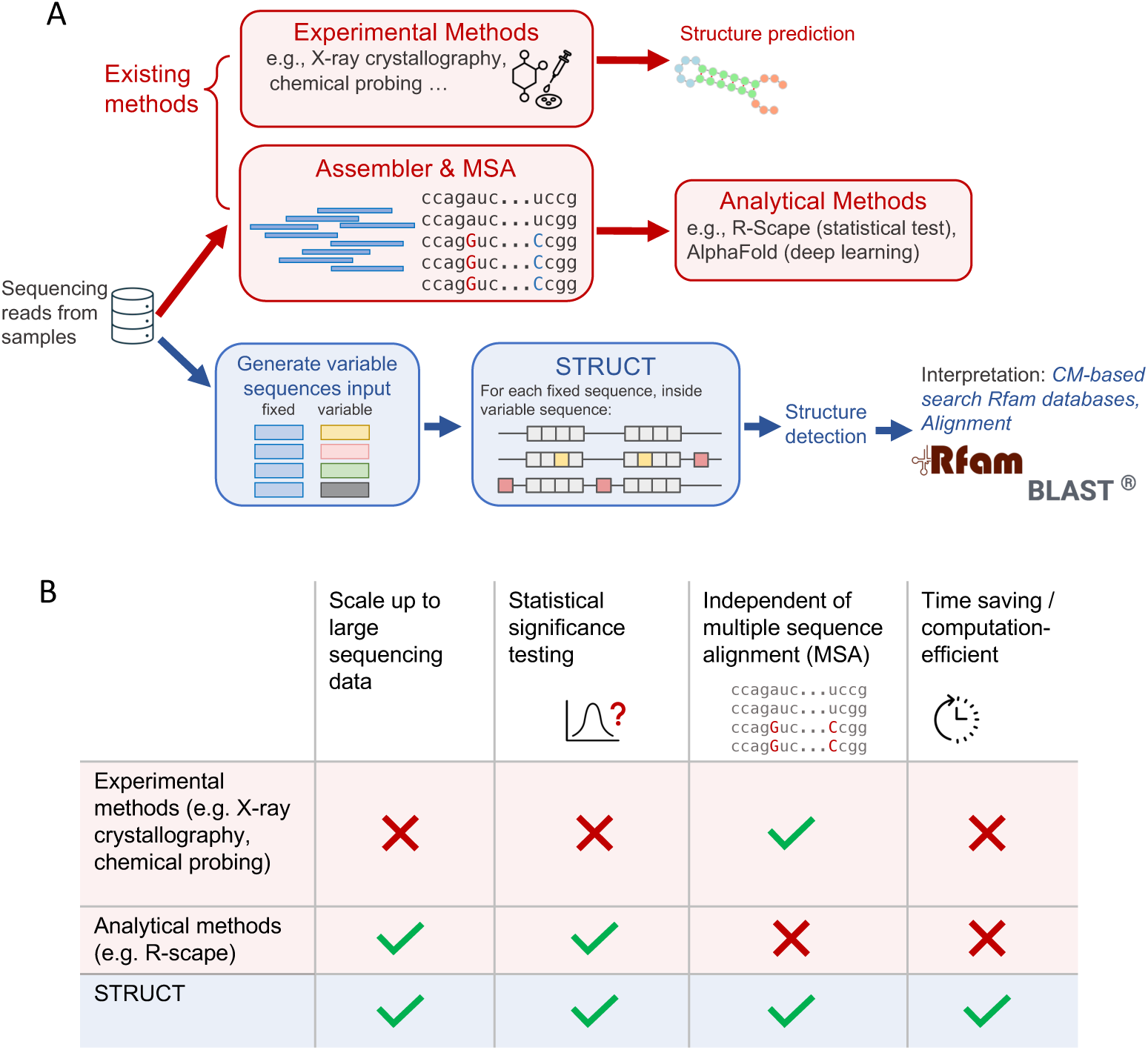
(A) Compared with existing experimental and analytical methods, STRUCT detects RNA structures by statistically analyzing variable sequences linked to a shared fixed sequence from raw RNA-seq data. Post-hoc interpretations include mapping to the Rfam database and BLAST search to the nucleotide database. (B) STRUCT is a high-throughput and efficient tool for detecting RNA structures without relying on assembly and MSA.

We first show that STRUCT can rediscover known structural elements on a purely statistical basis without relying on sequence alignment. We ran STRUCT on longitudinal HIV sampling from an HIV-infected cohort [23] on a host-by-host basis to control for confounding covariation caused by phylogenetic relations [11]. STRUCT recovers well-known structures without a reference sequence, including the ribosomal frameshift site, psi packaging signal, and trans-activating response element [24]. The power of STRUCT should be realized in its ability to predict conserved RNA structures missing from reference genomes. To test this, we ran STRUCT on the complex, multiorganism sequencing mixture from a mosquito metagenomics sample. Here, STRUCT prioritizes known structures in rRNA as a positive control while predicting structures in viruses where structural covariation has not been described and novel predictions in sequences missing from NCBI nucleotide databases. We thus present STRUCT as one of a growing number of tools [25] that predict structure from sequence.

Direct comparisons between STRUCT and existing covariation-based methods cannot be performed because STRUCT is the first statistical approach to detect RNA secondary structures without relying on MSA. However, we validated our structure predictions by comparing them with structures in the Rfam database and further interpreting them using nucleotide database query with BLAST.

## 2 Results

### 2.1 STRUCT is an MSA-free statistical approach to identify RNA secondary structures, bypassing MSA

STRUCT is a statistical approach designed to identify RNA secondary structures by focusing on conserved elements despite nucleotide variations at the primary level. We used SPLASH, a statistical algorithm to detect variations in raw sequencing reads, to generate input for STRUCT [26, 27]. We reasoned that if an RNA structure is functional, its overall structure would be conserved despite base changes in its nucleotide sequences, and sequences near this structure should be called by SPLASH. SPLASH processes raw sequencing data and generates analyzable units known as anchors and targets across the samples. An anchor is any specific k-mer sequence in a read, with each k-mer at a fixed offset downstream from said anchor the target. A given anchor may have multiple targets. STRUCT then detects patterns in target sequence variations across samples that could be explained by pressure to conserve secondary structures. The algorithm begins by selecting the most abundant target of an anchor as the base target, *T*_0_. It aims to identify a “stem-loop” configuration within *T*_0_ of each anchor (Figure 2A). Specifically, STRUCT looks for a consecutive k-mer (a minimum of 5 bases in practice) from the start of *T*_0_ and a reverse complement of this k-mer further downstream that does not overlap with the original k-mer. The reverse complement is defined according to Watson-Crick-Franklin base-pairing rules. The identified k-mer and its reverse complement are defined as the stems, and the intervening sequences are the loop. While RNA can fold following non-Watson-Crick-Franklin base-pairing rules [28], we only consider Watson-Crick-Franklin base-pairing to simplify the statistical formulation. As a result, our statistical tests tend to be conservative at identifying structures. Only anchors with a base target featuring a stem-loop structure are kept for further analysis. One limitation is that this approach primarily focuses on the simplest RNA secondary structure, stem-loops, and thus neglects more nuanced secondary structures such as bulges and pseudoknots.

**Fig. 2.**
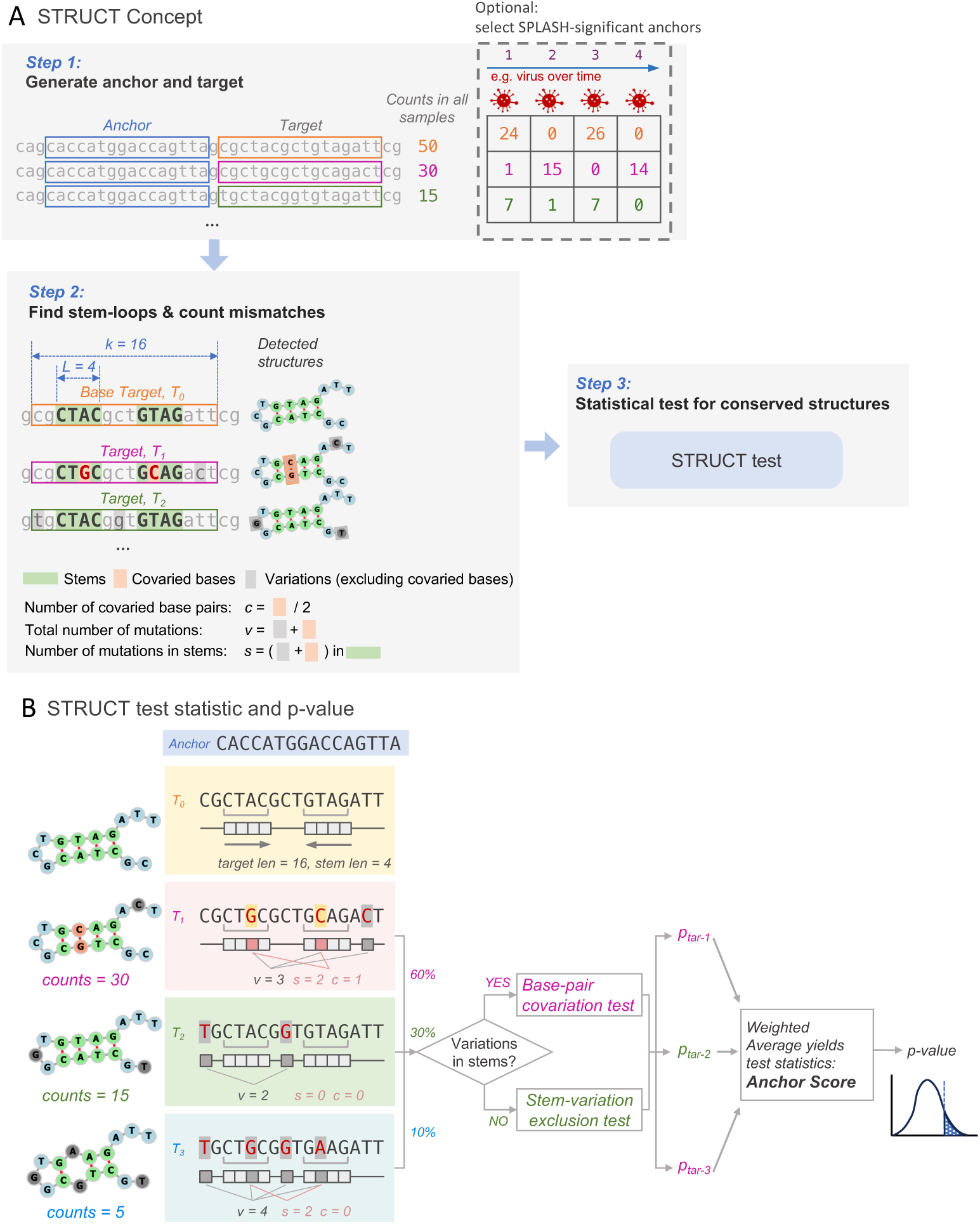
(A) STRUCT takes fixed sequences (anchors) and downstream variable sequences (targets) as inputs. In the toy example, an anchor of length 16 (highlighted in blue) is followed by multiple targets (highlighted in yellow, pink, and green) of the same length, offset by one base. In the study, we use SPLASH to generate anchors and targets and select SPLASH-significant anchors — target distributions of which differ across samples with statistical significance — for structure prediction. The second step involves identifying a stem-loop structure (illustrated with a *forna* RNA structure drawing) in the base target *T*_0_ and computing parameters (*c*, *v*, *s*) for the targets *T*_1_ and *T*_2_. These parameters are then used in the third step – statistical tests for detecting conserved structures (see Methods). STRUCT prioritizes conserved structures that exhibit sequence variation across different samples or over time. (B) Toy example of computation of STRUCT statistics. Stems in *T*_0_ are depicted as light gray boxes, with arrows indicating reverse complementarity. In the rest of the targets, STRUCT finds conserved base pairs (cartoon boxes in red) and tallies parameters *v*, *s*, and *c* (described in A). For each target, STRUCT tests the sequence for BPC or SVE based on the number of variations in stems (*s*). The test statistic, anchor score, is computed as the weighted average of these values, and STRUCT reports a closed-formed *p*-value.

STRUCT proceeds by assessing variation in the rest of the targets against the base target (Figure 2B). The algorithm tallies mismatches within or outside the stems and determines the compensatory mutations in stems. For instance, if a nucleotide A within stems of a base target changes to G in another target, and the corresponding T changes to C, this is considered a compensatory mutation pair that is consistent with a conserved structure, which we refer as compensatory mutations or base-pair covariation (BPC). Another mechanism consistent with conserved structures involves the absence of mismatches in the stems between targets despite their presence elsewhere. We call this mechanism stem-variation exclusion (SVE).

For each varied target, STRUCT combines the BPC and SVE tests. Each is tested against the null hypothesis that mismatches are uniformly distributed across any positions within the target and that these mismatches occur independently, given a fixed sequence length (*k*), stem length (*L*), and number of mismatches (*v*, Hamming distance from the base target). The target *p*-value, *p*_tar_ is defined for the two alternatives separately: 1) For BPC: *p*_tar_ is defined as the likelihood of having at least as many compensatory mutations as observed, given the predetermined parameters *k*, *L*, and *v*. 2) For SVE: *p*_tar_ is the likelihood of all mismatches occurring outside the stem regions. Both *p*_tar_ can be computed in closed form (Methods). The two target *p*-values are combined using an indicator function that checks for mismatches in the stem regions. If there are no mismatches, *p*_tar_ for SVE is used. Otherwise, *p*_tar_ for BPC is used.

Then, for each anchor, we construct an aggregate test statistic, anchor score, defined as the weighted mean of *p*_tar_ of all targets of an anchor. We can derive the distribution of the anchor score, compute a *p*-value for each anchor (Methods), and adjust *p*-values to control the false discovery rate (FDR) through the Benjamin-Hochberg (BH) procedure. With this hypothesis test, a low BH-adjusted *p*-value associated with an anchor suggests a stem-loop structure consistent with BPC or SVE among varied targets in samples.

We specified the method described so far as STRUCT (target mode), as we have been detecting structures in target sequences. We note that a stem-loop can be located distant from an anchor. Furthermore, natural stem-loop structures can vary substantially in length, and many structural elements possess greater complexities than simple stem-loops, such as internal loop. To comprise longer sequences and complexities, we adopted a similar analytical framework on compactors, i.e., STRUCT (compactor mode) (Methods; Suppl. Figure 1A). Compactors are longer local anchor-seeded assemblies sequences generated using probabilistic methods, and each seed-anchor can have multiple compactors (Suppl. Figure 1B) [29]. In this study, we generated compactors that are 81nt long. Anchors are included at the beginning as the first 27-mer of these compactors, and we call the 54-mer following the anchor the “trimmed compactor” (Figure 3A). The support of a compactor measures the count of occurrences in samples of all sequences it represents. The trimmed compactor of the compactor with the highest support is defined as the base trimmed compactor.

**Fig. 3.**
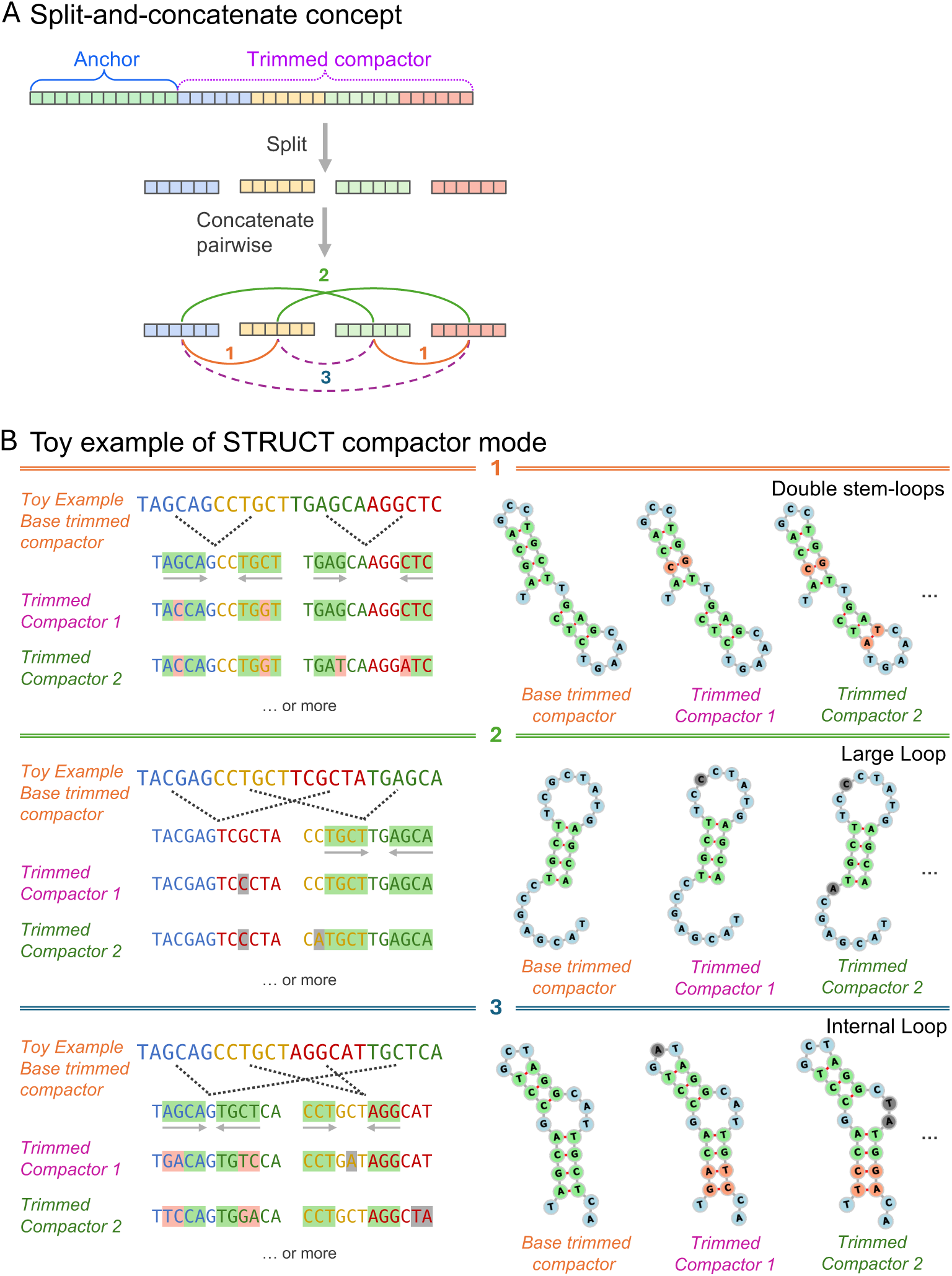
(A) STRUCT adopts a split-and-concatenate method for analyzing compactors. The process breaks a compactor into four distinct subsequences (colored blue, yellow, green, and red) and pairwise concatenates any two of these sub-sequences together. Three ways of concatenation are depicted in green, orange, and blue curves. (B) Toy example of 3 ways of split-and-concatenation of compactors. Each way results in two composite sequences. STRUCT aims to identify a stem-loop in each composite sequence. Thus, with three options of concatenations, compactors can be used to model more complex RNA structures: double stem-loops, stem-loops with large loop regions, or stem-loops with a bulge. forna RNA drawings of such structures are given for each case.

STRUCT (compactor mode) adopts a split-and-concatenate method for analyzing longer sequences, allowing for the identification of two pairs of complementary sequences in a compactor. The principle idea is to split a trimmed compactor into four nearly equal-length sub-sequences and then concatenate any two segments in an order-preserving manner (Figure 3A). This means that if one segment precedes another within the original compactor, it will also precede it in the new composite sequences. As a result, there are three ways to recombine four sub-sequences. In the study, we split a 54nt long trimmed compactor into 14, 14, 13, and 13 nt long sub-sequences, and the resulting recombined sequences have 28, 27, or 26 nucleotides (Methods). To detect structure, we treat each resulting recombined sequence like a target, allowing previously described processes in “target mode” to be directly applied to compactors. But now, because sequences are permuted, nested stems can be detected. This includes two adjacent stem-loops, a stem-loop with a longer loop, and a longer stem-loop with an internal loop and, in principle, pseudoknots (Figure 3B, Suppl. Figure 2). Next, for each split-and-concatenate permutation, the same statistical measure is applied to evaluate an anchor, with parameters being the aggregate of the two recombined sequences. This split-and-concatenate method can be expanded by dividing a long sequence into more than four sub-sequences. This increases the number of possible ways to reassemble the sequences, allowing for the detection of structures with more possibilities in future work (Methods). As we show in later sections, STRUCT (compactor mode) is more flexible and powerful than the target mode for detecting stem-loop structures. Therefore, we will primarily focus on reporting results from the compactor mode.

We designate anchors with a BH-adjusted *p*-value below 0.05 in target mode or compactor mode of STRUCT as structure-significant anchors. This distinguishes them from SPLASH-significant anchors, which are identified by the SPLASH algorithm as having significant variations across different samples. In the remainder of this paper, “significant anchors” refers to structure-significant anchors unless explicitly specified as SPLASH-significant anchors.

### 2.2 STRUCT *de novo* rediscovers conserved HIV-1 structural RNA elements

We tested STRUCT on human immunodeficiency virus (HIV) sequences. HIV is a single-stranded, positive-sense RNA virus characterized by a small genome, high mutation rates, and many secondary structures crucial to viral replication and host defense mechanisms [24]. These RNA structural motifs play critical regulatory roles, including promoting transcription or translation and direct virion assembly (dimerization and packaging) [24, 30, 31]. In HIV, the wide variety of known structural motifs and the presence of short stem-loops with considerable nucleotide variation make it an ideal case for STRUCT to detect RNA structures independently of traditional assembly and MSA techniques.

We analyzed publicly available Illumina short-read sequencing datasets of provirus sequences from a longitudinal study involving individuals infected with HIV-1 subtype B+ [23]. We applied STRUCT separately on seven hosts (Hosts 1124, 1194, 1211, 1532, 548, 583, and 746) to detect evolutionary conserved RNA structures, testing for significance within a host and controlling for confounding covariation caused by phylogenetic relations (Rivas, Clements, and Eddy 2017). STRUCT (target mode) identified 13, 5, 10, 7, and 3 significant anchors (BH-adjusted *p*-values ¡ 0.05) in Host 1124, Host 1194, Host 1211, Host 583, and Host 746, respectively, with no significant anchors detected in the other hosts. In Host 1124, all significant anchors match the HIV ribosomal frameshift signal element (RF00480) of the Rfam database, and the structural element is detected to be conserved through SVE. We show a detected example of a frameshift element in Figure 4A. Base-pairing within predicted stem regions shows consistency across various targets. Significant anchors detected in other hosts align to HIV genomes by BLASTn but do not match any structure in the Rfam database.

**Fig. 4.**
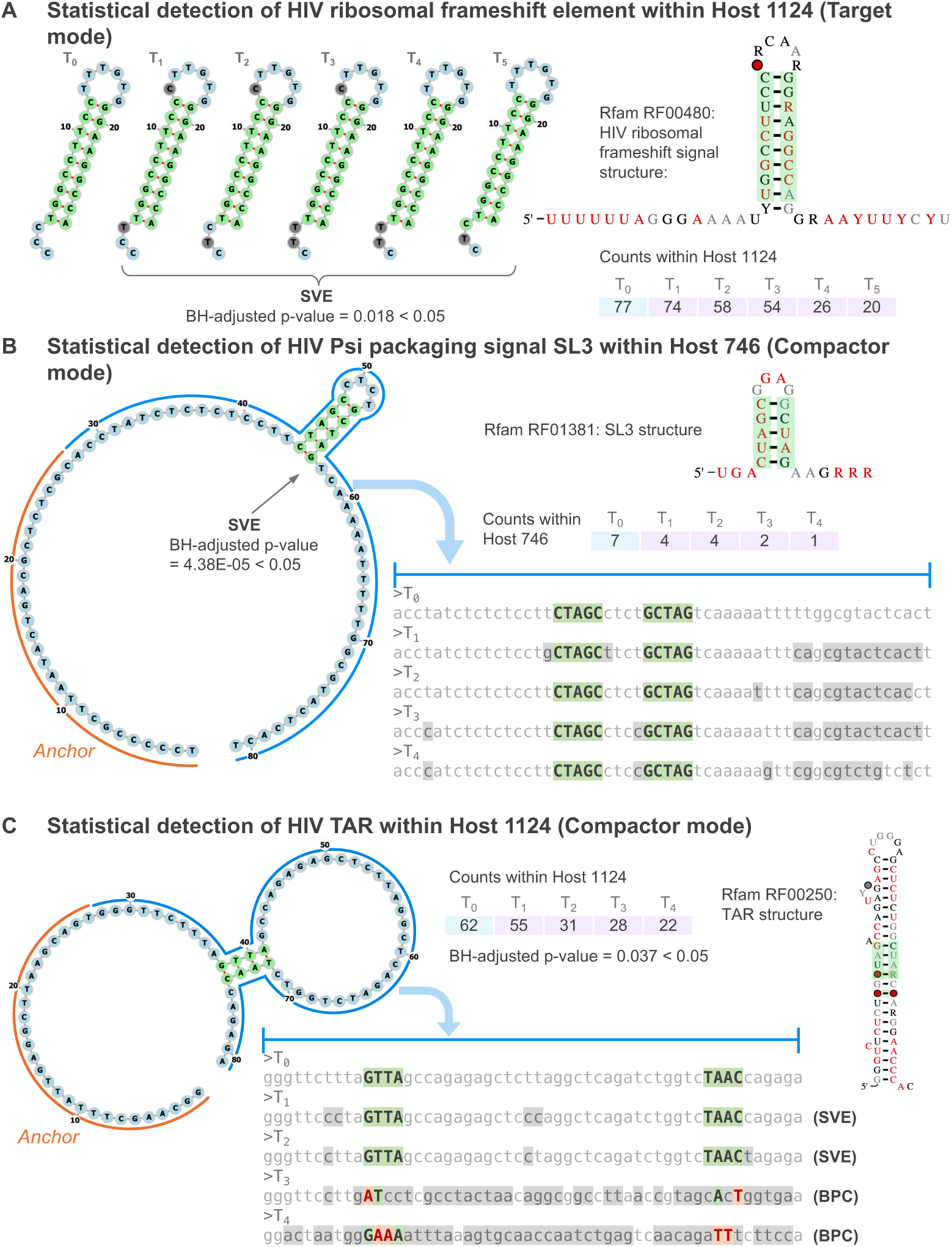
(A) – (C) show *forna* drawings of the detected structures that match HIV ribosomal frameshift element, HIV Psi packaging signal SL3 and HIV TAR, respectively, accompanied with the secondary structure in Rfam, *p*-value and counts of targets or compactors in samples. In *forna* drawings, nucleotides within the stems are green, and those outside stems are blue. (A) An example of the HIV frameshift element detected by STRUCT (target mode), conserved through SVE. The leftmost structure drawing is the base target. The rest to the right of the base target are varied targets. (B) An example of the HIV Psi packaging signal SL3 detected by STRUCT (compactor mode), conserved through SVE. The *forna* drawing is the base compactor. The aligned nucleotide sequences of varied compactor sequences are displayed. Capital letters and green highlights denote unvaried nucleotides in stems. Contrived bases are in red text color with orange highlight, and all other variations are shaded in gray. (C) Similar to (B), an example of the HIV TAR detected by STRUCT (compactor mode), conserved through SVE in *T*_1_ and *T*_2_, and BPC in *T*_3_ and *T*_4_.

STRUCT (target mode) is expected to overlook elements due to the algorithm’s design, which disregards sequences with minimal variations across samples, consequently missing structures in highly conserved regions. Additionally, the target mode may miss structural elements for the following reasons: 1) Targets (here 27 bps) are too short to accommodate structural elements with longer stems or larger loops; 2) False negative calls of anchors in SPLASH (Methods), which we considered tangential to this study and did not investigate in depth. To investigate 1), we concentrate on reporting results for STRUCT (compactor mode).

STRUCT (compactor mode) identifies a median of 2.2% structure-significant anchors relative to the total number of SPLASH-significant anchors equipped with compactors across all hosts. We found hits to the retroviral psi packaging element stem-loop 3 (SL3; RF01381) in 4 out of 7 hosts (except for Host 1194, 548, and 1532), one of the most frequently detected. SL3 is a 14 nt highly conserved stem-loop, and STRUCT (compactor mode) precisely identifies the stem sequences conserved through SVE in compactors (Figure 4B). STRUCT detected sequence variations outside the stem-loops, particularly on both sides of the stem-loop, making it less possible to be identified near an anchor in the target mode.

STRUCT (compactor mode) also detects the trans-activation response element (TAR; RF00250) in 4 out of 7 hosts. The TAR has a hairpin structure essential for viral replication that contains 24 base pairs, a loop of 6 bases, and three bulges in the stems. The nearest bulge to the loop is four nucleotides away (Figure 4C). In target mode, the STRUCT cannot detect the stem-loop due to its requirement for a minimum stem length of 5 nucleotides. Consequently, the bulge must be at least five nucleotides away from the loop for target mode to identify the stem-loop successfully. Compactor mode has the advantage of detecting compensatory stems that are further apart and away from the loop, making detection of TAR possible. Overall, we observe that STRUCT (compactor mode) exhibits greater ability to identify conserved structures; therefore, we will focus on the results from the compactor mode.

### 2.3 STRUCT identifies conserved rRNA structures in mosquito metatranscriptomics

Mosquitoes are vectors and hosts for various microorganisms, including eukaryotic, prokaryotic, and viral agents. Metatranscriptomic sequencing of individual mosquitoes offers a single assay capable of sampling from diverse organisms, many likely lacking references. We sought to identify functionally important elements within complex metatranscriptomics data by prioritizing regions under selection for conserved structural elements in order.

We analyzed a publicly-available mosquito metatranscriptomics dataset that profiled 148 diverse wild-caught mosquitoes (Adult *Aedes*, *Culex*, and *Culiseta* mosquito species) collected in California [32]. STRUCT (target mode) identified 687 significant anchors in all reads, and STRUCT (compactor mode) identified 4718 significant anchors. We searched for matches of the combined anchor and base target sequences and base compactors in the Rfam database. Our findings revealed that under compactor mode, base compactors of 190 significant anchors match ribosomal RNA (rRNA) in Rfam, a positive control for STRUCT, because rRNA has highly conserved structures across diverse organisms despite variations in primary sequences. Among the top 10 most significant anchors (lowest *p*-values) that have their base compactors matched to rRNA, two matched with eukaryotic large subunit rRNA (28S rRNA), four matched the eukaryotic small subunit rRNA (18S rRNA), two matched bacterial large subunit rRNA (23S rRNA), and two matched bacterial small subunit rRNA (16S rRNA).

Figure 5A shows a match to 16S rRNA. Within the base compactor, two stemloop structures of stem lengths of 5 and 6, respectively, were detected. BPCs occur at different locations in the stems, and these changes can vary across compactors. For example, the base pair G-C (nucleotides at index 52 and 64) in the base compactor has a compensatory change to A-T in the first compactor followed but remains unchanged in the next compactor and only has one base change in the last one. The base pair A-T (nucleotides at index 48 and 68) in the base compactor changes to T-A in all three compactors. Similarly, STRUCT observed covariations of base pairs in the compactors that match 28S rRNA (Figure 5B).

**Fig. 5.**
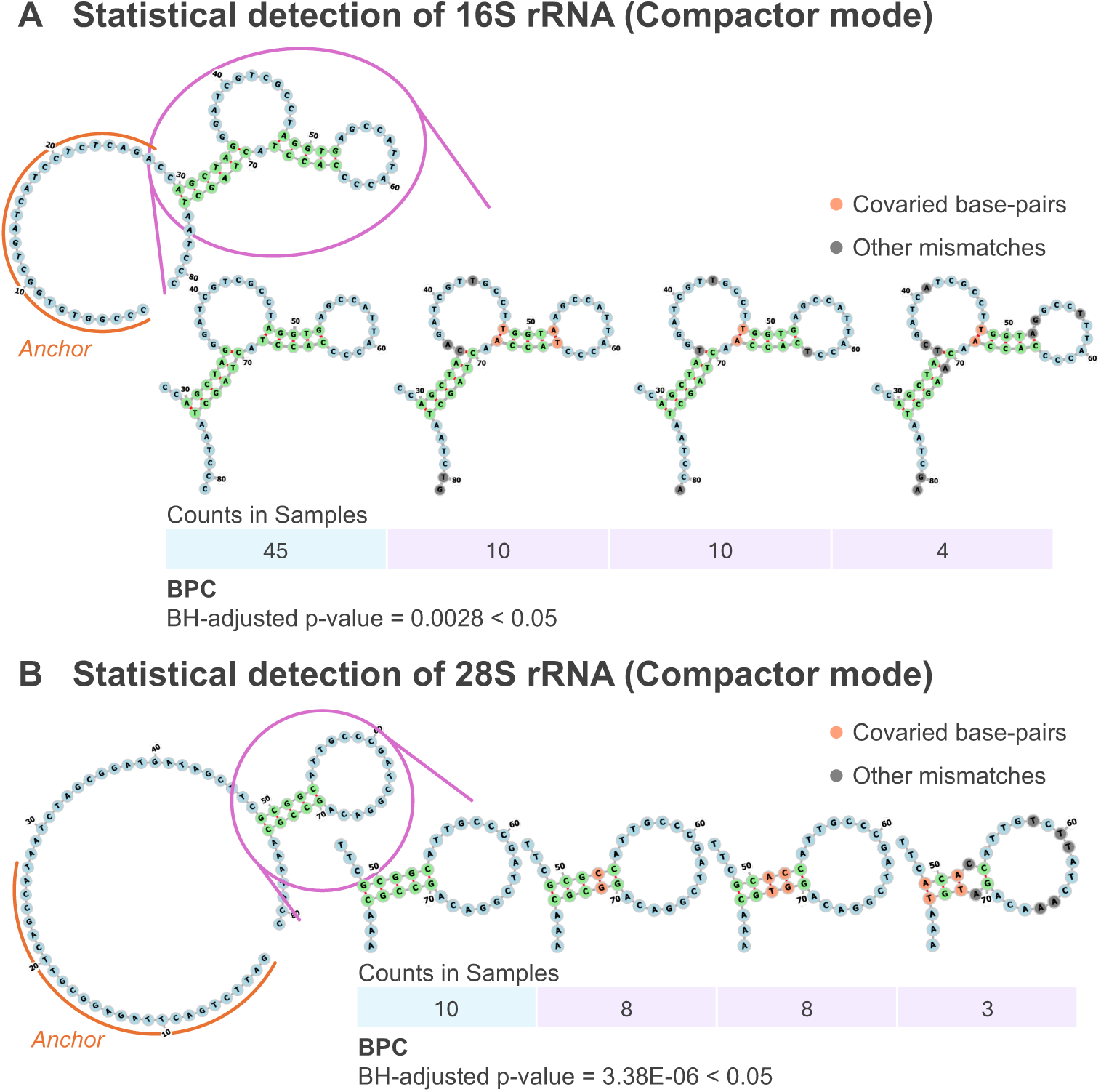
(A) An example of the detected structures that match 23S rRNA by STRUCT (compactor mode), conserved through both SVE and BPC across targets. (B) An example of the detected structures that match 28S rRNA by STRUCT (compactor mode) conserved through BPC. The zoom-ins show the contrived base pairs in the varied sequence of the detected stem-loop.

### 2.4 STRUCT identifies RNA viral structures in mosquito metatranscriptomics

4113/4718 (87.18%) base compactors aligned (BLASTn) to Arthropda phylum genomes, 55/4718 (1.17%) to bacteria, 51/4718 (1.08%) to chordata phylum, 46/4718 (0.96%) to viruses, 28/4718 (0.59%) to fungi, and other organisms (Figure 6A), with 391/4718 (8.28%) having unaligned base compactors. Given the myriad functions of RNA structures in regulating virus replication, we first focused on sequences that align with viruses in the BLAST results.

**Fig. 6.**
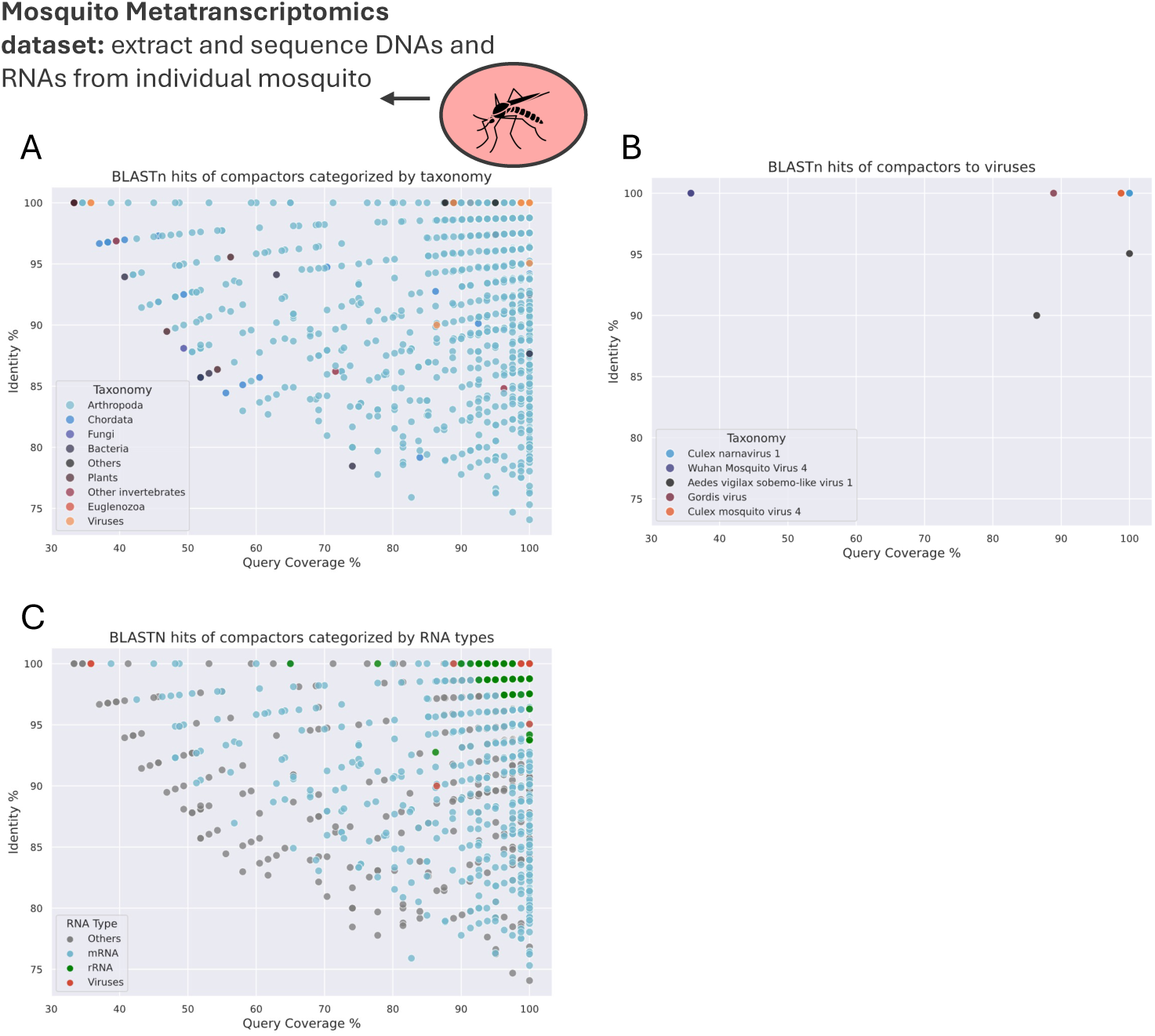
(A) Each point represents a unique structure-significant base compactor detected by STRUCT (compactor mode), with colors indicating taxonomy obtained from the BLASTn hit with the lowest E-value. The plot does not show the relative abundance of compactors across samples. Query coverage refers to the percentage of the query sequence aligned with the subject sequence. Percentage identity indicates the proportion of identical nucleotides in the aligned region. (B) Each point represents a distinct structure-significant base compactor aligned to viral genomes using BLASTn. These points correspond to the viral taxonomy group “Viruses” shown in Figure A. Sequence that aligned with Wuhan Mosquito Virus 4 has low coverage but 100% similarity. (C) Each point represents a unique structure-significant base compactor under STRUCT (compactor mode), as in the previous plots, but points are categorized by RNA type. BLASTn hits with the smallest E-value are used for labeling. Query coverage and percentage identity are defined as in the previous plot.

STRUCT detects five viruses without relying on assembly of reads and provides statistical evidence of potentially functional elements within them: Culex narnavirus 1, Gordis virus, Culex mosquito virus 4, Aedes vigilax sobemo-like virus 1, and Wuhan mosquito virus 4. Notably, Aedes vigilax sobemo-like virus 1 and Wuhan mosquito virus 4 were not reported in previous analyses, highlighting STRUCT’s ability to detect RNA viruses without references. Culex narnavirus 1 had the highest number of aligned base compactors (40 out of 46). For all viruses except Wuhan mosquito virus 4, the base compactors aligned with over 80% coverage and high sequence similarity. The base compactors aligned to Wuhan mosquito virus 4 showed only 29 out of 81 bases with 100% sequence similarity to the subject sequence (Figure 6B). We queried other compactors of the same anchor; two of the four aligned with Wuhan mosquito virus 4, and exhibit 71% query coverage with near-perfect sequence identity. The unaligned sequences potentially originate from the same virus; however, the absence of a complete reference genome or additional sequencing reads precludes definitive identification. We hypothesize that similar instances of incomplete detection of less abundant viruses are more pervasive than expected in metatranscriptomic datasets.

We show evidence of BPC at various positions in predicted stem-loops of Culex narnavirus 1. In Figure 7A, the G-C base pair (nucleotides at indices 46 and 51) in the base compactor changes to A-T in 3 out of 4 subsequent compactors. The only compactor without BPC at these positions exhibits another pair of compensatory changes – A-T to G-C at indices 44 and 53. The extensive BPCs also make it possible to detect structures in target mode (Figure 7B). We also show examples of the statistical detection of structures of Gordis virus and Culex mosquito virus 4 by SVE (Figures 7C and 7D). Although the detected structures are located near the anchor (within 27 bases downstream), the lack of sequence variations causes them to be ignored in target mode. Variations occurring at the ends of the compactor sequences directed these sequences to be analyzed under compactor mode.

**Fig. 7.**
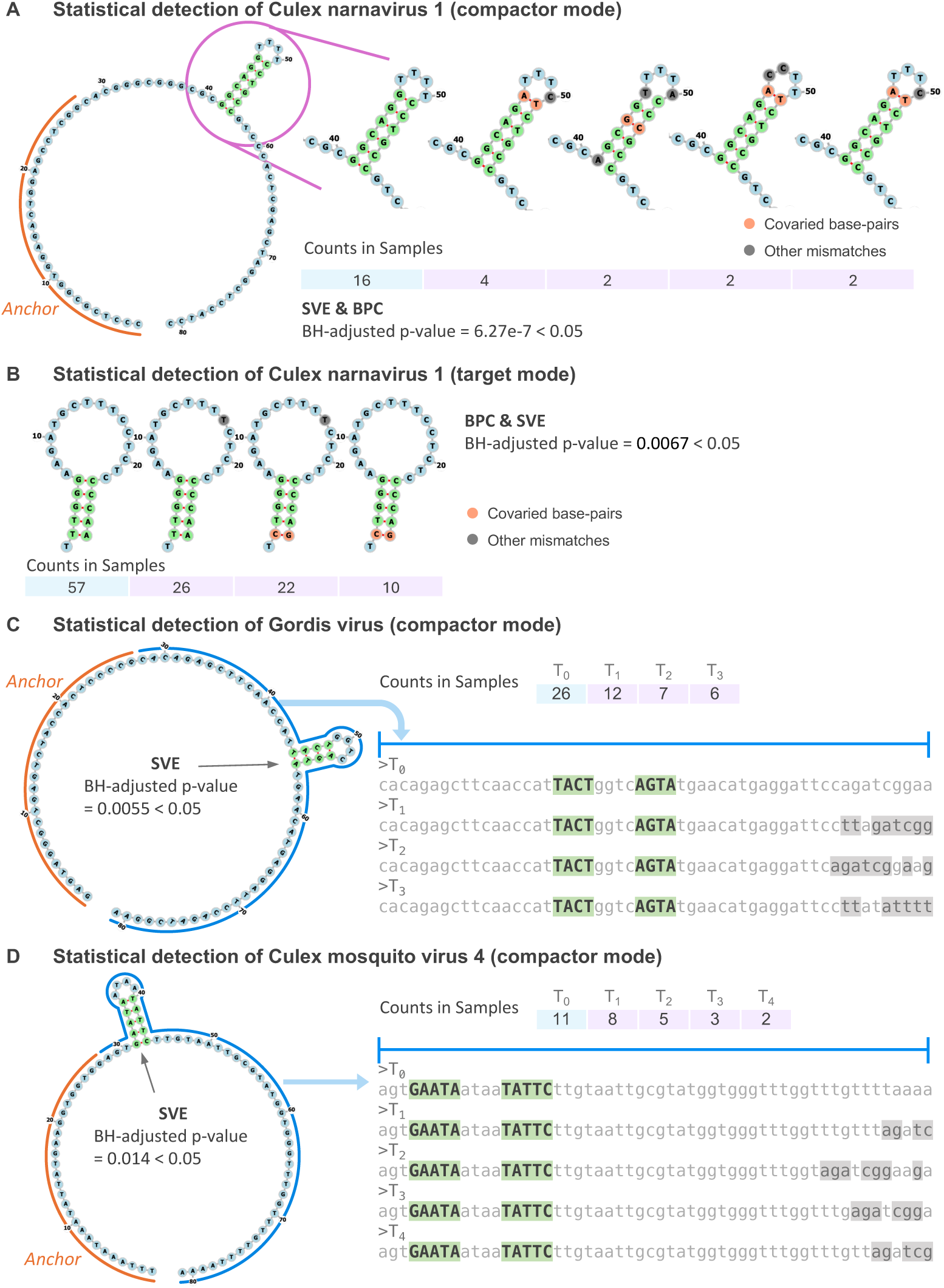
(A) – (B) show *forna* drawings of detected structures that are mapped to the Culex narnavirus 1, separately identified under compactor mode and target mode of STRUCT, accompanied with *p*-value and counts of targets or compactors in samples. The structure detected in compactor mode is conserved through BPC. The structure detected in target mode is conserved through SVE in *T* 1 and BPC in *T* 2 and *T* 3. (C) STRUCT (compactor mode) detected Gordis virus via structure conserved through SVE. The *forna* drawing is the base compactor. The aligned nucleotide sequences of varied compactor sequences are displayed. Capital letters and green highlights denote unvaried nucleotides in stems. Conserved bases are in red text color with orange highlight, and all other variations are shaded in gray. We chose to show a stem with a length of 4 nucleotides and a tetraloop, rather than the 5-nucleotide stem with a 2-nucleotide loop detected by the algorithm, because tetraloop sequences are more stable and avoid the increased strain from the sharp bend in a 2-nucleotide loop. (D) STRUCT (compactor mode) detected Culex mosquito virus 4 via structure conserved through SVE. The color scheme convention is the same as in (C).

### 2.5 STRUCT detects eukaryotic mRNA structures with statistical evidence of BPC or SVE across species in mosquito metatranscriptomics

Eukaryotic mRNAs have structure that is important for their function [33, 34]. Because STRUCT weights stem-loop covariation, and the genes encoding eukaryotic mRNAs are not expected to rapidly accumulate mutations (structurally compensatory or otherwise), we did not generally expect it to detect mRNA structures unless multiple related species separated by large evolutionary distances were present. Interestingly, the Batson et al. dataset provided sequences from 10 *Aedes*, *Culex*, and *Culiseta* related mosquito species [32]. After running STRUCT, 1132 out of 4718 (23.99%) base compactors of the significant anchors had BLASTn aligned to mRNA (Figure 6C; Methods). This finding supports the hypothesis that eukaryotic mRNAs with statistical evidence of BPC or SVE can be identified using STRUCT when applied to datasets that contain mRNA sequences from multiple related but sufficiently divergent species.

### 2.6 STRUCT detects novel sequences by statistically evident structures with BPC or SVE in mosquito metatranscriptomics

Unaligned (by BLASTn) base compactors suggest the presence of novel sequences not currently documented in the existing nucleotide database. STRUCT identified them with statistically evident RNA structures. To investigate and interpret these unexplored sequences, we generated longer compactors (over 81 nucleotides) for the significant unaligned anchors (see Methods). In total, 669 compactors were created, with lengths ranging from 135 to 459 nucleotides (Suppl. Figure 3). Re-running BLASTn with these extended compactors led to the identification of sequences that aligned with Culex Bunyavirus 2 and Culex narnavirus 1 (Figures 8A and 8B). This suggests that the previously unannotated anchors of these compactors may correspond to viral genomes that have not yet been documented, highlighting STRUCT’s ability to identify novel viral sequences with statistically evident structures.

**Fig. 8.**
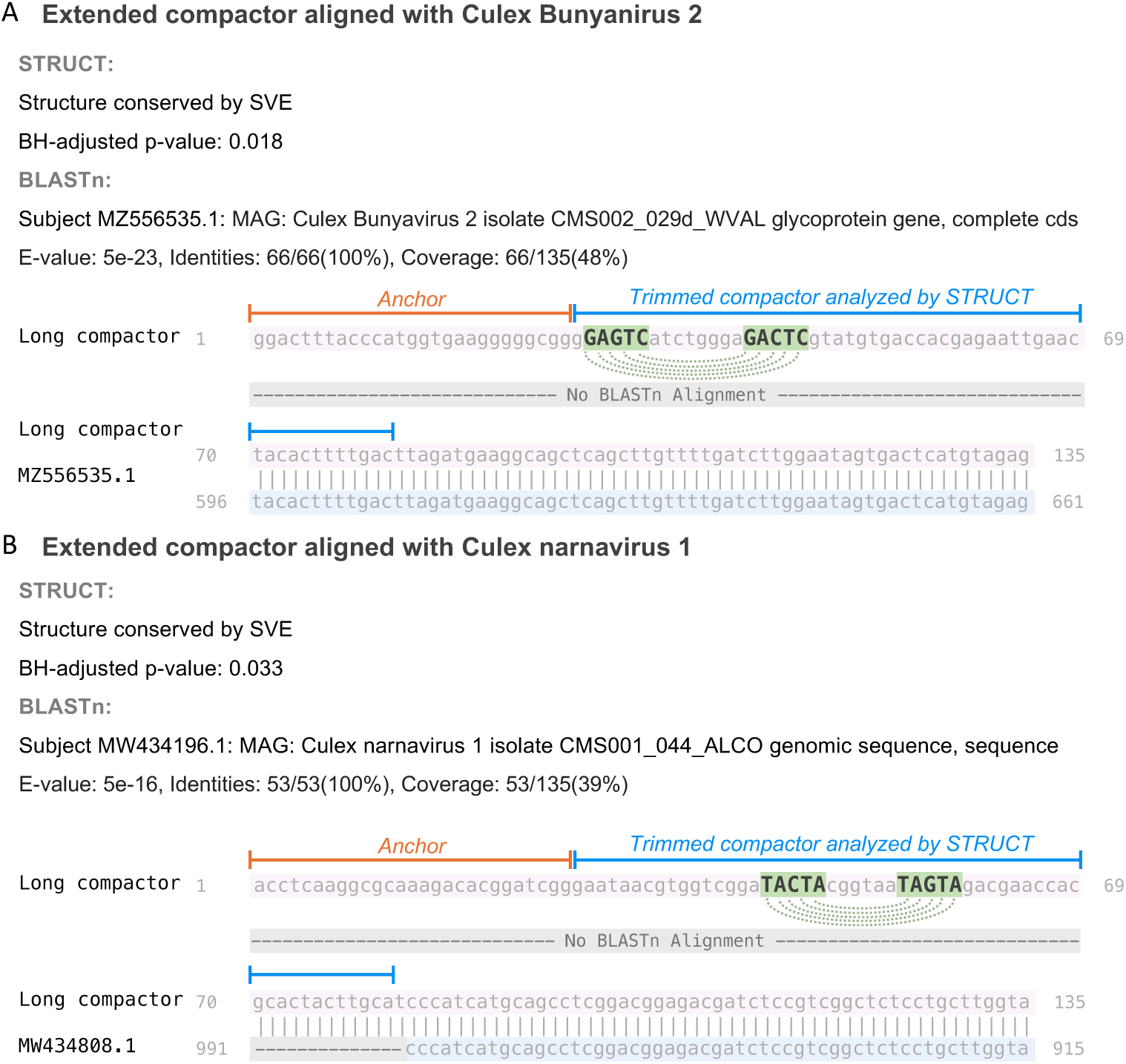
(A) An extended compactor of 135 nt of an unaligned 81nt compactor by BLASTn aligns with a subject sequence of Culex Bunyanirus 2. The alignment between the subject sequence (highlighted in blue) and the extended long compactor (highlighted in purple) is shown, along with BLASTn E-value, identities, and coverage. The anchor sequence, marked with orange whiskers, and the compactor where STRUCT searches for stem-loops, marked in blue, are indicated. Nucleotides in the detected stems are colored green and shown in uppercase. (B) Similar to (A), using the same color scheme and marker notation, an extended compactor of 135 nt from an unaligned 81nt compactor aligns with a subject sequence from Culex narnavirus 1.

For compactors that remained unaligned after the second round of BLASTn, we performed *in silico* translation and queried protein databases using BLASTx. We found sequences whose translated proteins matched uncharacterized proteins at various *loci* in mosquitos. This suggests that STRUCT can potentially uncover novel sequences linked to proteins.

## 3 Discussion

Amid an explosion in available RNA-sequencing data, and an ongoing need to understand the RNA structurome across the tree of life, scalable sequence-based RNA structure prediction algorithms are needed. We developed a method for RNA structure prediction, STRUCT, that takes variable sequences (targets) linked to a shared fixed sequence (anchors), generated from raw sequencing data in a reference-free manner, and interrogates these sequences for predicted stem-loop RNA structures. We combined two statistical tests on these sequence pools: a base-pair covariation (BPC) test and a stem variation exclusion (SVE) test that appear to preserve predicted structures. As a result, biologically meaningful RNA structures are more likely to be identified. Unlike existing sequence-based RNA structure prediction approaches, STRUCT does not rely on multiple sequence alignment (MSA) or assembly and can identify RNA structures from a single unknown sequencing sample. Furthermore, though structure prediction might be more accurately performed with long-read sequencing data, since most publicly available data consists of short reads, STRUCT is advantageous as it effectively predicts structure from short-read datasets and provides critical statistics to address whether predicted structures are conserved.

We tested STRUCT in isolated HIV-1 provirus sequencing and mosquito meta-transcriptome datasets without providing any information other than short-read sequencing data. From HIV amplicons generated from patients’ blood, STRUCT identified several known HIV genome RNA secondary structures, including the frameshift signal, the psi packaging signal, and the TAR. STRUCT did not pick up all of the predicted or known structures of HIV as we only applied STRUCT to SPLASH-significant anchors, but the top hits of STRUCT were all RNA structures necessary for the HIV life cycle.

In RNA-sequenced whole mosquitoes, we found that STRUCT identified ribosomal subunit RNAs. STRUCT also identified structured RNA sequences that, when queried against known sequences, were found to belong to RNA viruses. This finding highlights the unique power of STRUCT in identifying RNA viruses in mixed or unknown sequencing samples. Furthermore, our algorithm identified several sequences that align with viruses not identified in the Batson et al. analysis [32]. Moreover, some longer query sequences from the unaligned sequences generated by the compactor assembly function of SPLASH are incompletely aligned to known viral species. Intriguingly, these could represent related virus sequences that have not yet been added to the NCBI reference sequence database and remain to be worked up. Indeed, reference genomes for Aedes vigilax sobemo-like virus 1 and Wuhan mosquito virus 4 became available after the published analysis by Batson et al., which may have prevented their identification.

Despite STRUCT’s effectiveness in identifying known structured RNAs and generating hypotheses about new regions of RNA structure, we recognize the following limitations in its current implementation:

1. Bulges and conserved bulges: STRUCT views sequencing data as anchor-target pairs to explore various target sequences linked to the same anchor. The method identifies the perfect reverse complement sequence within the most abundant target (base target) and counts several types of mismatches observed in different targets associated with the same anchor. This approach, primarily focused on the most straightforward RNA secondary structures such as stem-loops, may neglect the potential presence of bulges in double-helical structures.
2. Watson-Crick-Franklin base-pairing: In the probability formulations, STRUCT adopts canonical Watson-Crick-Franklin base-pairing to identify potential stemloops. While essential, Watson-Crick-Franklin base-pairing often receives excessive focus, overshadowing other crucial interactions in RNA structure, such as non-Watson-Crick-Franklin pairings that are vital for RNA three-dimensional structure and function. The overemphasis on Watson-Crick-Franklin base-pairing and the underemphasis on non-base-pairing interactions in predicting RNA structures can lead to an incomplete understanding of RNA’s structural and functional capabilities [28].
3. Choice of sample: STRUCT is best suited to sequencing data derived from organisms with structured RNA and rapidly changing RNA, which emphasizes its use in detecting RNA viruses and suggesting structured elements in an unknown mixture. For example, though human polyadenylated mRNAs have secondary structure, mutations in the genomic DNA template occur very slowly, making compensatory mutations in stem-loop structures unlikely. Although STRUCT identified some mRNA stem-loops when applied to a metatranscriptomics dataset that contains mRNA sequences from related but sufficiently divergent mosquito species, we therefore recommend STRUCT as best utilized in settings where the source RNA sequences are hypothesized to include some variation, ideally A) the presence of known or unknown RNA viruses or viroids, or B) DNA-templated RNA from related organisms across evolutionary time (e.g., RNA-seq from related primates (or mosquitoes) or within a species separated by many generations).

We plan to address these limitations in future work. Ultimately, STRUCT conducts covariation analysis in sequences and provides statistical evidence for predicted structures, all in a reference sequence-agnostic and MSA-free manner. Doing so may help discover new biology hiding in sequencing datasets, especially the presence of RNA viruses, in which secondary structures play an outsized role in the life cycle. Downstream, after identification and validation, targeting of RNA structures (including viral structures) with small molecules is an emerging human disease therapeutic area [35]. We encourage using STRUCT on RNA sequencing datasets, whether derived from known viral RNA inputs or from complex mixtures of organisms.

## 4 Conclusion

STRUCT is an MSA-free, assembly-free, metadata-free statistical approach to detect RNA secondary structures from raw sequencing data, quantifying base-pair covariations or stem variation exclusion in the putative RNA structures. STRUCT rediscovers known HIV structural elements in pro-virus sequences and identifies rRNA structure in metatranscriptomics data. STRUCT also finds viral structures in metatranscriptomics, showing potential for viral disvovery in mixtures of organisms. The results point to STRUCT as a useful tool for identifying structures in RNA-seq datasets, opening the door to rapid RNA structure prediction and hypothesis generation.

## 5 Methods

### 5.1 Generating data input for STRUCT using SPLASH

The SPLASH algorithm was applied to the raw FASTQ sequences of each dataset to generate analyzable units known as anchors and targets (both 27 nt long) across the samples. [26, 27]. An anchor is any specific k-mer sequence in a read, with each k-mer at a fixed offset downstream from said anchor the target. A given anchor may have multiple targets.

SPLASH version 2.1.15 was run with the following command, taking samplesheet.csv as an argument, which stores paths to FASTQ sequences:

`$ splash –-outname_prefix result –-anchor_len 27 –-gap_len 0`

`--target_len 27 –-poly_ACGT_len 8 –-max_pval_opt_for_Cjs 0.1`

`--n_threads_stage_1 4 –-n_threads_stage_2 32 –-n_bins 128`

`--n_most_freq_targets 10 –-kmc_max_mem_GB 12 samplesheet.csv`

We select SPLASH-significant anchors and their targets as input for STRUCT. The output file `{results.after_correction.scores.tsv` contains such anchors and their top 10 targets, specified by `–-n_most_freq_targets 10`.

Note that SPLASH is chosen as a tool to generate inputs for STRUCT. The derivation of the SPLASH test statistic involves splitting data into train and test sets, which introduces inherent randomness and can lead to false negatives.

### 5.2 Generating compactors from SPLASH-significant anchors

Compactors were generated using software compactors, a local assembly method based on a probabilistic model included in SPLASH version 2.1.15 [29]. The following command extends seed anchors listed in `anchor_list.txt`, queries the raw FASTQ files specified in `samplesheet.csv`, and outputs the compactors for seed anchors and their support – the number of sequences that a compactor represents – to `compactors.tsv`. Compactors of 81 nucleotides in length are generated by specifying `--num_kmers 2 and –-max_length 81`.

`$ compactors –-kmer_len 27 –-num_kmers 2 –-epsilon 0.05 –-beta 0.4`

`--lower_bound 1 –-max_mismatch 4 –-poly_threshold 6 –-max_length 81`

`--max_anchor_compactors 10 –-num_threads 16 –-reads_buffer_gb 48`

`samplesheet.csv anchor_list.txt compactors.tsv`

In cases where we needed to extend the compactors to interpret the sequences, we adjusted the `–-max_length` flag to 1000 to generate longer compactors, capping their length at 1000 nucleotides.

### 5.3 Removing contaminant sequences

We implemented filtering procedures on the anchors to eliminate genomic artifacts and PCR primers, ensuring that STRUCT results were free from common sequencing contaminants. An internal pipeline was applied to efficiently index reference sequences and query every k-mer present in combined anchor and target sequences and compactors against these references. For each queried sequence, the output includes a comprehensive list of matching references of each k-mer. We set a threshold for the maximum allowable counts of a contaminant reference and filtered out sequences containing more hits than this threshold. Additionally, for the HIV datasets, sequences containing PCR primers used in a specialized PCR protocol were filtered out [36]. This filtering was accomplished using the command `grep –v` to exclude sequences matching the primers.

### 5.4 Filtering out low-abundance and low-variability sequences

As specified, up to the top 10 most abundant targets or compactors are generated for each anchor. Targets or compactors with less than 5% abundance relative to the generated sequences were discarded for better computation efficiency. Next, anchors with fewer than three varied targets or compactors, excluding the base target or base compactor, were also excluded from the analysis. Under compactor mode, we also filtered out anchors whose second most abundant compactor has only support 1, i.e., the sequence it represents only shows up once in the dataset. This step ensured only anchors with sufficient variability in their targets or compactors were considered. As a result, short stem-loops in highly conserved sequences may be intentionally overlooked under these settings.

### 5.5 STRUCT (target mode)

#### 5.5.1 Likelihood of observed sequence variations in a target: target ***p*-value, (*p*_tar_)**

Consider the most abundant target of an anchor the base target, *T*_0_. We search for a stem-loop structure with a minimum stem length of 5 in *T*_0_. The algorithm achieves this by selecting a 5-mer from the beginning of *T*_0_ and searches for its reverse complement from the base immediately following the selected 5-mer, adhering to the canonical Watson-Crick base pairing rules of A to T and C to G. If no reverse complement is found downstream, the algorithm steps one base forward and repeats the search for the next 5-mer. If a reverse complement is located, the algorithm extends the current k-mer and its reverse complement by one base and verifies if the extended sequences remain complementary. This recurring process halts when the extended sequences no longer match or when they begin to overlap, whichever occurs first.

The length of the stem is *L* (*L* ≥ 5). We then iterate all targets *T_i_* (*i* = 1, · · · *, K*, *K* is the number of targets of an anchor besides *T*_0_), and calculate the number of single-nucleotide mismatches between *T*_0_ and *T_i_*, denoted as Hamming distance *v*. We then compute how the mismatches are distributed in the stem-loop.

Under the null hypothesis, with fixed target length *k*, stem length *L*, and number of mismatches *v*, we assume that mismatches occur uniformly are uniformly distributed across any *v* positions out of *k* in the target *T_i_*. Furthermore, each mismatched base is independent of others and transitions to any of the other three bases with equal probability (for example, A changes to C, G, or T with equal likelihood).

The alternative hypothesis proposes that a stem-loop structure is preserved in two mechanisms. The first mechanism preserves the stem-loop structure through mismatches that form compensatory mutations within the stems, termed base-pair covariation (BPC). The second mechanism considers that mismatches are selectively excluded from the stems, called stem variation exclusion (SVE).

In the context of BPC, we define the conditional likelihood of having at least as many compensatory mutation pairs *C* as the observed value *c*, given the predetermined parameters target length *k*, stem length *L*, and number of mismatches *v*, as the first target *p*-value, denoted as 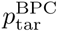,

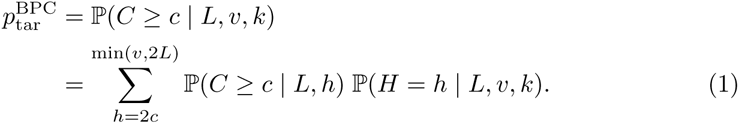

where the last line is allowed by the law of total probability. *H* denotes the variable representing the number of mismatches observed in stems, and *h* denotes the possible values that *H* can take on.

In the preceding equation, the second term can be modeled with a hypergeometric distribution,

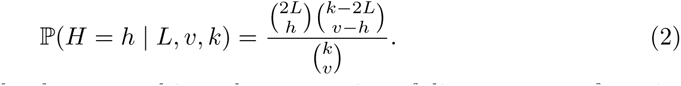

The first term can be decomposed into the summation of discrete cases of *g* pairs of compensatory mutations existent in a target, and it can be further decomposed and computed in a closed form (see Supplementary Methods).

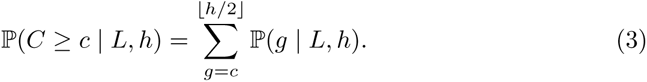

To have statistical power against another alternative, where mutations are absent in stem regions (SVE), we define another statistic, 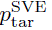, representing the probability of *v* mismatches all occurring outside stems,

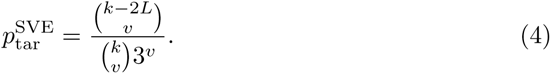

The final likelihood, denoted *p*_tar_, combines the two alternative hypotheses of the preservation mechanisms of stem-loops via an indicator function. The observed number of mismatches in stems, *s*, serves to distinguish between the two mutually exclusive cases: BPC (*s >* 0) and SVE (*s* = 0). Accordingly, we employ 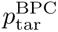 in cases where mismatches are present in stems, 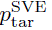 when mismatches are absent in stems. Formally, *p*_tar_ is given by

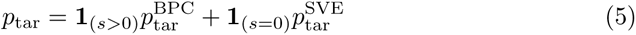

which follows an approximate uniform distribution in a discrete sense. Given this uniformity, the exact distribution of *p*_tar_ can be computed as there is a limited number of possible combinations for the locations and types of mutations in a target.

#### 5.5.2 Test statistic, anchor score *R*

To capture the likelihood of preserving a stem-loop structure among varied targets, we assign a single statistic to an anchor, termed anchor score, *R*. We define *R* as a linear combination of each target *p*_tar_ weighted by their abundance. Formally, consider an anchor with *K* targets *T*_1_, *…*, *T_K_*. Define the weight of the *i*-th target, *w_i_* (where *i* = 1*, …, K*), as the proportion of counts of that target among all *K* targets. Represent the weights of the *K* targets with the vector ***w*** = (*w*_1_*, …, w_K_*)^⊤^ with 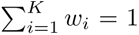, and the corresponding *p*_tar_ as 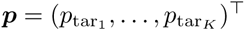. Then, the anchor score *R* is

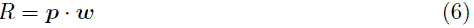

#### 5.5.3 *p*-value

To obtain a *p*-value for the test that measures the global conservation of stem-loops, we need the null distribution of the test statistic, anchor score, *R*. Since *R* is the weighted mean of *K* discrete uniformly distributed random variables, *p*_tar_, the distribution of random variable *R* can be derived using convolution of discrete variables. For example, *K* = 2, we expand *R*,

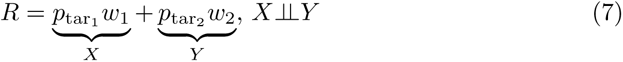

where *w*_1_ and *w*_2_ are constants and *p*_tar1_ and *p*_tar2_ are PMF of known distributions. Define two independent random variables, *X* = *p*_tar1_ *w*_1_ and *Y* = *p*_tar2_ *w*_2_, supported on *R_X_*and *R_Y_* respectively, with PMF *p_X_* (*x*) and *p_Y_* (*y*). The distribution of anchor score *S* can then be derived via convolutions of the two independent random variables, *X* and *Y*, i.e.,

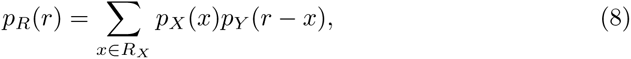

where *r* is the observed value of the anchor score variable *R*. Now a *p*-value is computed as the lower-tail probability,

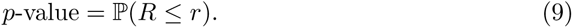

Since we conduct a hypothesis test and compute a *p*-value for each anchor, to control the false discovery rate (FDR), we apply the Benjamini-Hochberg procedure over the list of *p*-values from the multiple hypothesis tests. We report the list of corrected *p*-value through Python functionality `statsmodels.stats.multitest.multipletests` with FDR controlled at *α* = .05.

### 5.6 Simulation of anchor score, *R*

To study the distribution of test statistic (anchor score, *R*) across multiple samples under the null hypothesis and to validate the BH-adjusted *p*-value computation, we performed simulations of anchor score under null and compared the empirical cumulative density function (ECDF) of *p*-values with the empirical data.

The simulation procedure is as follows. Given a pre-processed dataset containing anchors, each anchor has a base target *T*_0_ (a stem-loop was detected in *T*_0_), and targets *T_i_* (*i* = 1*, …, K*). For each anchor, we have also computed targets weights *w* = (*w*_1_*, …, w_K_*)^⊤^ and tallied the number of mismatches in each target *v* = (*v*_1_*, …, v_K_*)^⊤^. Subsequently, for each anchor, we simulated *K* targets. The *i*th simulated target was generated by randomly selecting *v_i_* positions in the base target, *T*_0_, and replacing the nucleotide at those positions with one of the other three nucleotide types, each with equal probability. We then computed a conditional likelihood for each simulated target given fixed stem length *L*, number of mismatches *v*, and target length *k*, same as the computation of *p*_tar_. Next, we performed the same process to compute the simulated test statistic, anchor score *R*, and *p*-value for each anchor in the given dataset (Suppl. Figure 4A).

We conducted this simulation on the pre-processed mosquito metatranscriptomics dataset, repeating the process 1000 times. The empirical cumulative distribution function (ECDF) of the simulated *p*-values was then plotted and compared with the empirical data (Suppl. Figure 4B).

### 5.7 STRUCT (compactor mode)

The compactor mode of STRUCT operates on compactors generated from SPLASH-significant anchors. Anchors are included as the first 27-mer of these compactors, and the 54-mers that follow are referred to as trimmed compactors. Define the trimmed compactor of the most supported compactor as the base trimmed compactor.

#### 5.7.1 Split-and-concatenate method to process compactors

STRUCT adopts a split-and-concatenate method to process trimmed compactors and incorporate longer and more complex structures in our predictions. We split a 54nt long trimmed compactor into four nearly equal-length sub-sequences of 14, 14, 13 and 13 nt long, following index slicing: [0: 14), [14: 28), [28: 41), [41: 54). These sub-sequences are designated as *D*^(1)^, *D*^(2)^, *D*^(3)^ and *D*^(4)^. The sub-sequences are then concatenated into two recombined sequences in an order-preserving manner, ensuring that sub-sequences with smaller indices precede those with larger indices. As a result, we have the following three ways to obtain two recombined sequences:

- *D*^(1)^ − *D*^(2)^: 28nt, *D*^(3)^ − *D*^(4)^: 26nt
- *D*^(1)^ − *D*^(3)^: 27nt, *D*^(2)^ − *D*^(4)^: 27nt
- *D*^(1)^ − *D*^(4)^: 27nt, *D*^(2)^ − *D*^(3)^: 27nt

Such split-and-concatenate methods can be expanded by dividing a long sequence into more than four sub-sequences. This division increases the number of possible ways to reassemble the sequences, allowing for the detection of structures of more possibilities. For example, we split a trimmed compactor into eight nearly equal-length sub-sequences. Concatenating any two sub-sequences with their orders preserved will result in 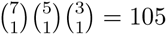 combinations.

#### 5.7.2 Computing anchor score under compactor mode

Under each result of split-and-concatenate, each trimmed compactor turns into two recombined sequences, as described in the previous section, and we try to find a stem-loop within each of them. If no stem-loop is found in any of the two, this split– and-concatenate way is discarded for the currently evaluated anchor.

We treat compactors under each split-and-concatenate way as targets, allowing all established algorithms for the target mode to be directly applied to compactors. To map the treated compactors to our target framework, we separately compare the two recombined sequences of a trimmed compactor to their corresponding indices in the base trimmed compactor and compute the parameters necessary computing *p*_tar_, including stem length *L_j_*, number of mismatches *v_j_*, number of compensatory mutation pairs *c_j_*, and number of mutations in the stem regions *s_j_*, where *j* denote the specific recombined sequence (*j* = 1 or 2).

Next, we compute the likelihood of the observed variations in compactors as *p*_tar_, using the aggregated parameters: total stem length (*L* = *L*_1_ + *L*_2_), total number of mismatches (*v* = *v*_1_ + *v*_2_), number of compensatory mutation pairs (*c* = *c*_1_ + *c*_2_), number of mutations in the stem regions (*s* = *s*_1_ + *s*_2_), and the combined lengths of k-mers (*k* = *k*_1_ + *k*_2_ = 54). Finally, anchor score (*R*) and *p*-value for compactor mode are computed analogously as they are for targets.

### 5.8 *p*-value distribution from datasets

We ran STRUCT on a mosquito metatranscriptomics dataset and seven HIV Hosts’ provirus sequencing data (Table 1). The empirical distributions of STRUCT BH-adjusted *p*-values of anchors validate that STRUCT assigns significance to sequences containing conserved stem-loops, either through BPC or SVE (Suppl. Figure 5). It also validates that STRUCT effectively identifies significant sequences of structural RNA, including rRNA or RNA viruses, within real sequencing data.

**Table 1.** Datasets and software.

| RESOURCE | SOURCE | IDENTIFIER |
| --- | --- | --- |
| **Public data** |  |  |
| HIV-1 | Antar et al. [23] | BioProject accession: PRJNA613107 |
| PCR primers for HIV-1 provirus extraction | Bruner et al. [36] |  |
| Mosquito metatranscriptomics | Batson et al. [32] | BioProject accession: PRJNA605178 |
| Rfam database | Kalvari et al. [37] | https://rfam.org/ |
| **Software and algorithms** |  |  |
| STRUCT | This study |  |
| SPLASH & SPLASH2 (version 2.1.15) | Chaung et al. [26],<br>Kokot et al. [27] | https://github.com/refresh-bio/SPLASH |
| Compactors | Henderson et al. [29] | https://github.com/refresh-bio/SPLASH |
| BLAST | Camacho et al. [38] | https://blast.ncbi.nlm.nih.gov/Blast.cgi |
| Infernal (version 1.1.4) | E. P. Nawrocki and S. R. Eddy [39] | http://eddylab.org/infernal/ |
| forna | Kerpedjiev et al. [40] | http://rna.tbi.univie.ac.at/forna/ |

### 5.9 Datasets and software included in the study

This study analyzes existing publicly available data, listed in Table 1 with accession numbers. Software and algorithms used in this study are also provided in Table 1.

### 5.10 Query of structure match in the Rfam database

We conducted a query of structure matches to the Rfam database for compactors (from compactor mode) and combined anchor and target sequences (from target mode) from the HIV and mosquito metatranscriptomics datasets. The sequences were saved to FASTA files. The search utilized the software Infernal (version 1.1.4), which implements covariance modes (CMs) to search for similar RNA structures in the Rfam database.

`$ cmscan –Z $TOTAL_RESIDUAL –-notextw –-rfam –-cut_ga –nohmmonly`

`--cpu 32 –-tblout <OUTPUT_PATH> –-fmt 2 –-clanin Rfam.clanin Rfam.cm`

`<FASTA_PATH>`

### 5.11 Exploratory sequence querying strategy for mosquito metatranscriptomics dataset

We ran remote BLASTn to query base compactors of structure-significant anchors against the nucleotide database nt. The queried sequences were saved to FASTA files. BLASTn (remote) was run using the following command (E-value threshold: 1E-3, word size: 11):

`$ blastn –remote –db nt –query <FASTA_PATH> –evalue 1e-3 –out`

`<OUTPUT_PATH> –task blastn –outfmt 6 –max_target_seqs 4 –dust no`

`-word_size 11 <FASTA_PATH>`

The sequence hit is chosen based on the lowest E-value. Species are determined using the hit’s accession number and categorized based on their superkingdom (for bacteria and viruses) or phylum (for eukaryotes). The categories include arthropoda, bacteria, chordata, viruses, fungi, plants, other invertebrates, euglenozoa, and others. BLASTn hits were also classified by RNA – rRNA, mRNA, RNA virus, and others – according to the subject sequence title to which a query sequence aligns.

For base compactors that did not return any BLASTn hit from the previous query, we generated longer compactors (length longer than 81 nts) for these anchors. In the resultant outputs, compactors of length 81 were excluded, and the rest were named long compactors. We then did a second round of sequence querying against the nt database for long compactors using BLASTn. For sequences that still did not have any hits, we ran BLASTx, which translates the nucleotide query sequence in six reading frames, resulting in six protein sequences. These translated sequences were searched against the nr database, with combined statistics across the six frames to assess statistical significance. The BLASTx command:

`$ blastx –remote –db nr –query <FASTA_PATH> –evalue 1e-3 –out`

`<OUTPUT_PATH> –task blastx –outfmt 6 –max_target_seqs 6 –word_size 6`

### Declarations

- Data availability: This study analyzes existing publicly available data, listed in Table 1 with accession numbers.
- Ethics approval and consent to participate: Not applicable.
- Competing interests: No competing interest is declared.
- Author contribution: J.F.W., A.R., and J.S. conceived and designed the study, J.F.W. and J.S. designed the statistics, J.F.W. performed analyses, J.F.W., A.R., and J.S. interpreted results and wrote the manuscript.

## Supplementary Information

1. Supplementary Methods.
2. Supplementary Figures.
3. Supplementary Tables. This file contains lists of structure –significant anchors that have Rfam or BLASTn annotations, detected by STRUCT (compactor mode) output.

## Supplementary Figures

**Suppl. Figure 1:**
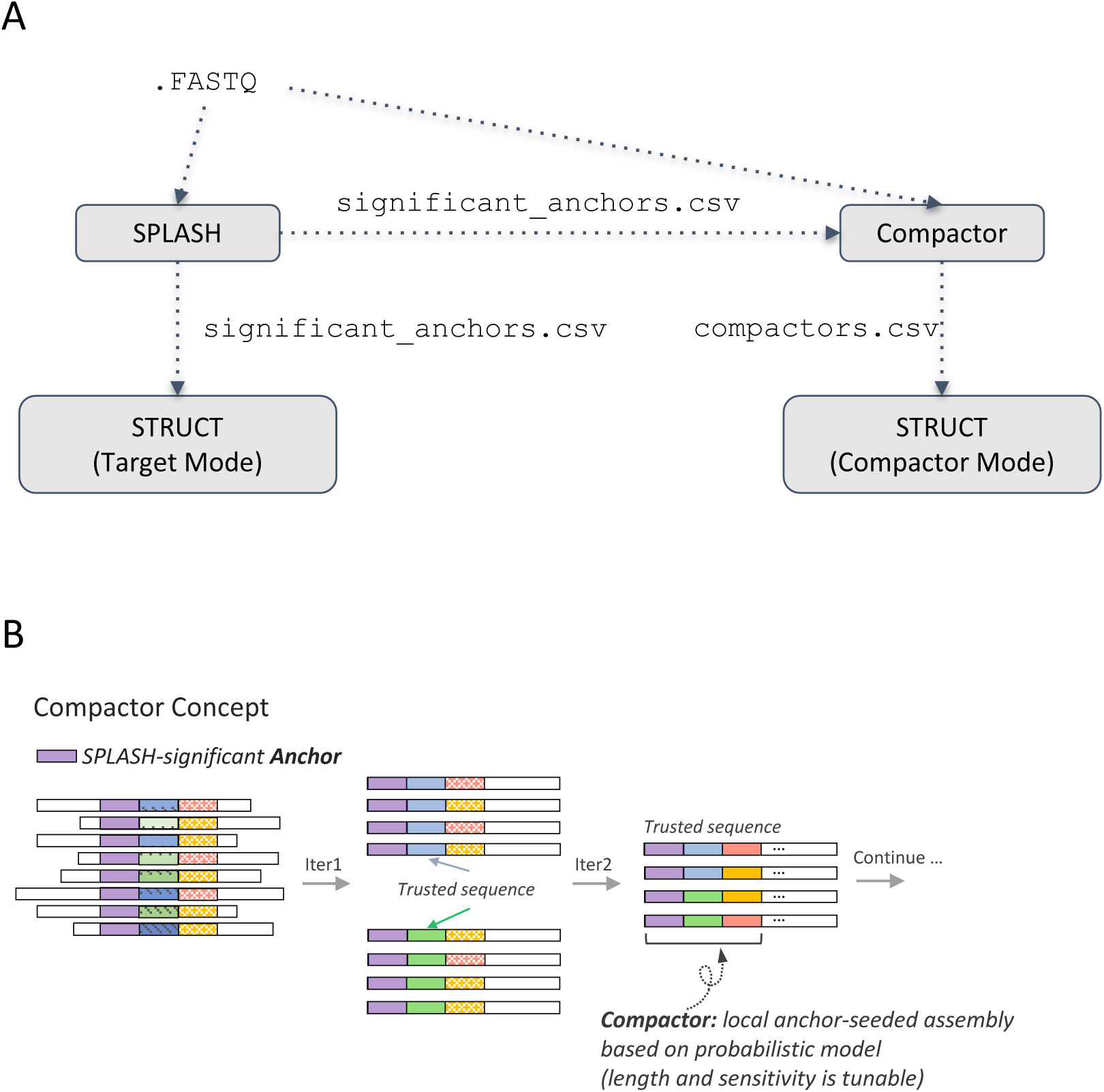
STRUCT (compactor mode) workflow and compactor generations. (A) The raw sequencing data in FASTQ format is initially analyzed by SPLASH, generating an output file significant anchors.csv that contains the SPLASH-significant anchors and their targets. Compactors are then constructed using the SPLASH-significant anchors as seeds to query the original FASTQ file, and deposited in the output file compactors.csv. The targets and compactors derived from the SPLASH-significant anchors are subsequently used as inputs for the two modes of STRUCT: Target Mode and Compactor Mode, respectively. Dotted lines represent file dependencies between the processes. (B) Compactors are local anchor-seeded assemblies of regions that vary across samples. They are generated iteratively based on a probabilistic model.

**Suppl. Figure 2:**
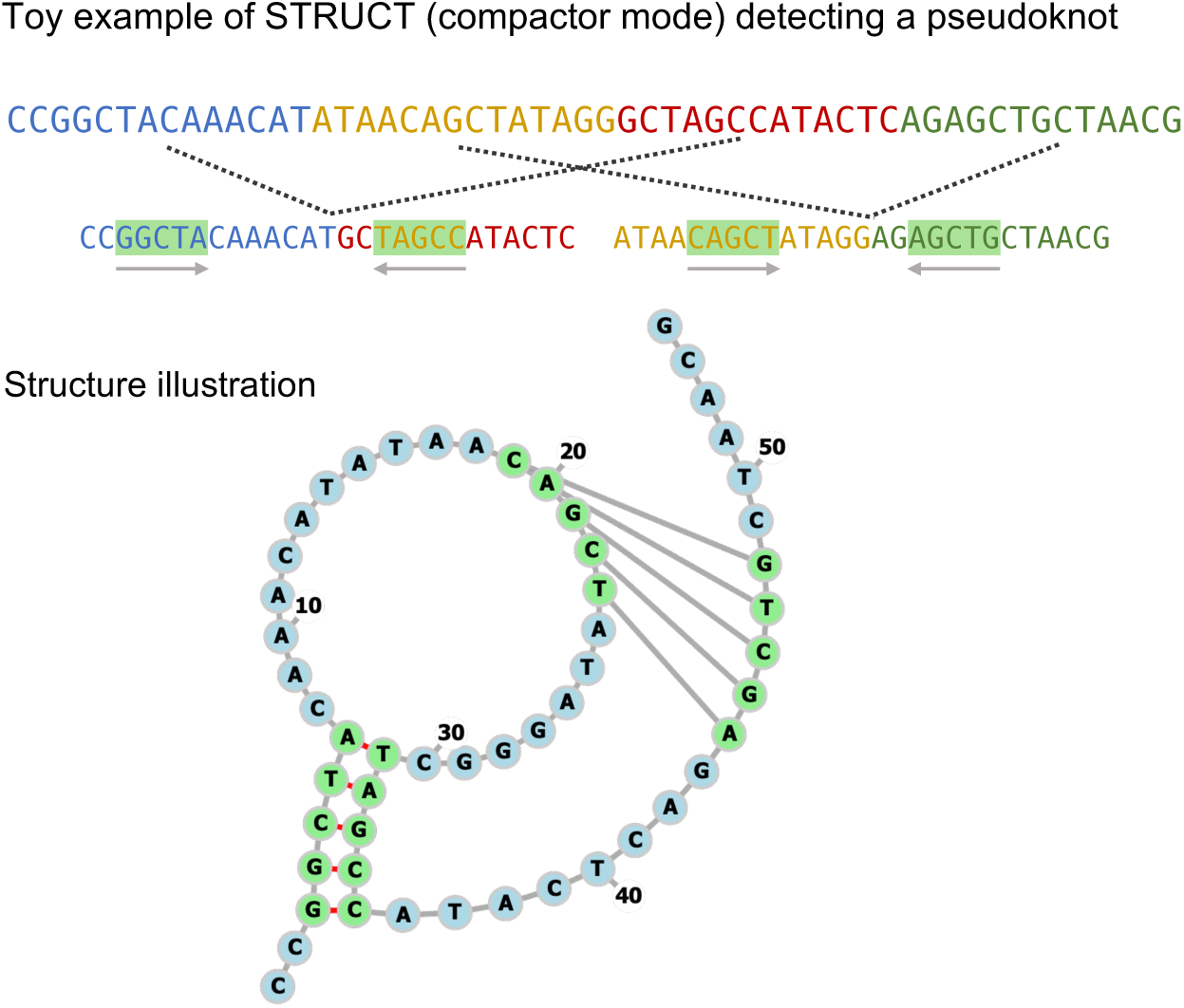
Toy example of STRUCT (compactor mode) detecting a pseudoknot. In principle, STRUCT can detect pseudoknots in sequences. The toy example demonstrates that a trimmed compactor adopts the second method of split-and-concatenation, as illustrated in Figure 3. This results in two composite sequences, with compensatory stems identified in both. A forna drawing of the detected pseudoknot is provided, with nucleotides in stems highlighted in green circles and pseudoknot links represented by gray straight lines.

**Suppl. Figure 3:**
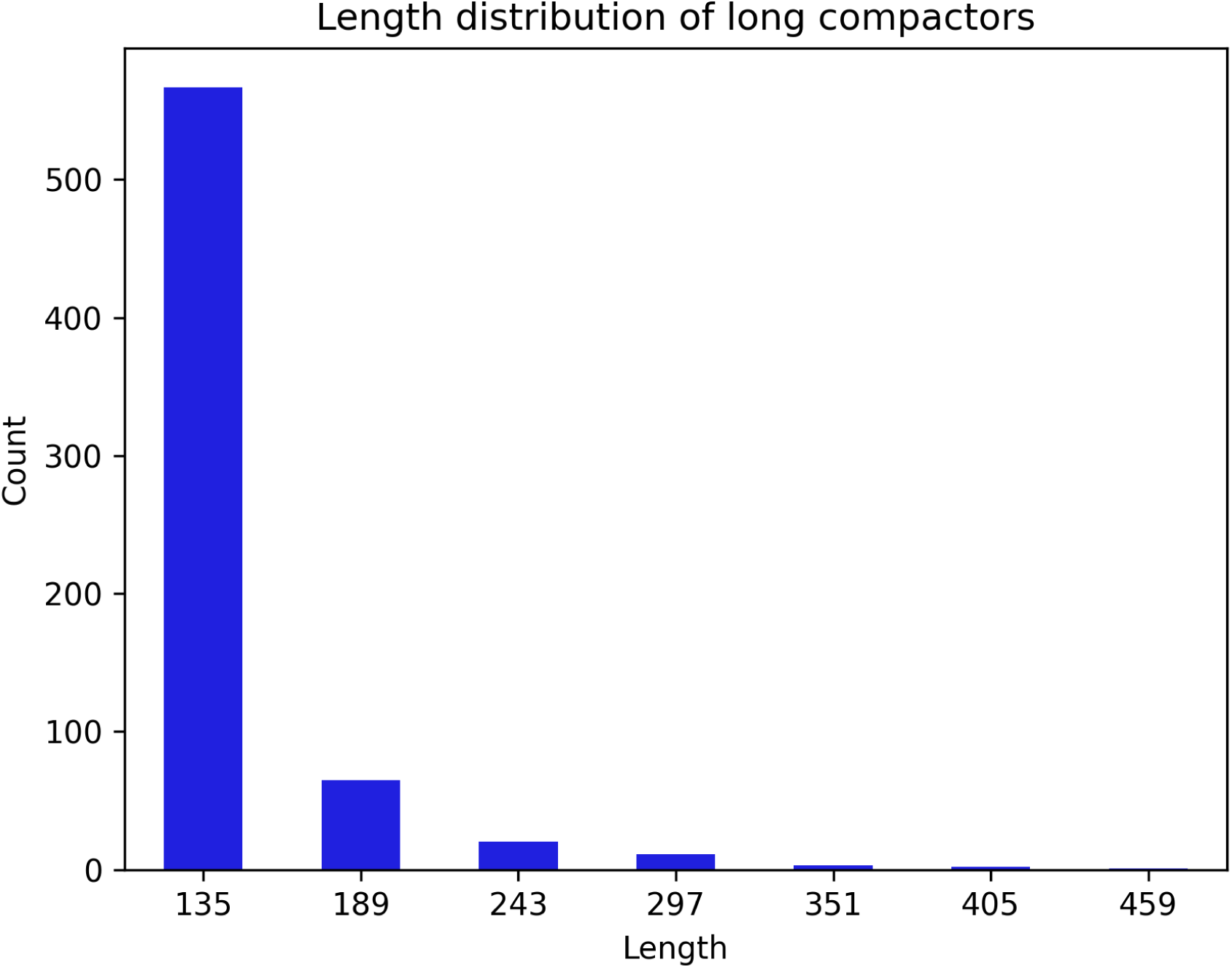
Length distribution of extended compactors over 81 nucleotides (nts). To interpret unannotated base compactors (81 nt) using BLASTn, longer compactors with lengths capped at 1000 nucleotides are generated for the anchors of these base compactors. The length distribution of these compactors is visualized in the bar plot.

**Suppl. Figure 4:**
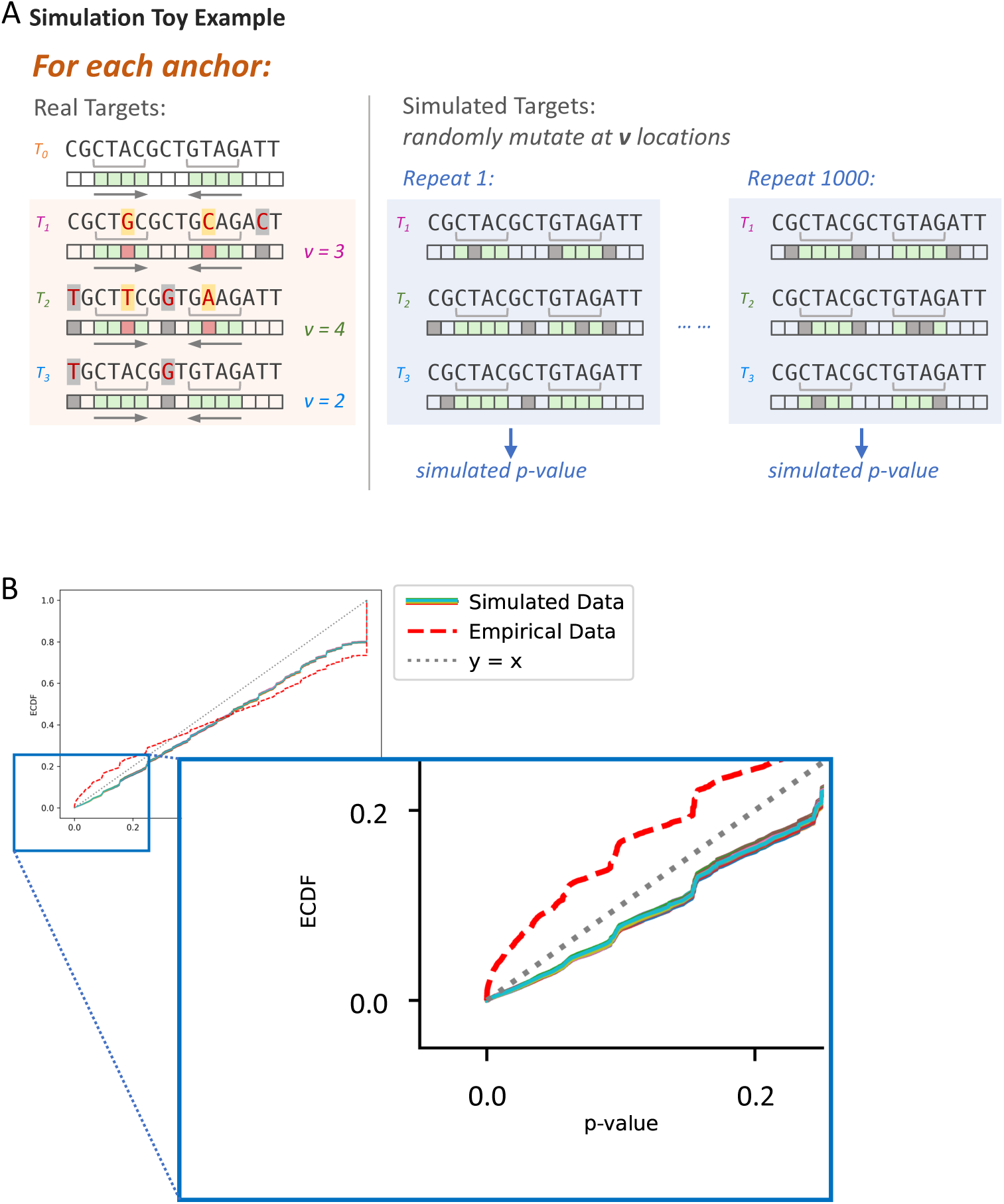
Comparison of *p*-value distribution of STRUCT ran on empirical and simulated data. (A) A toy example illustrating the simulation procedure of test statistic, anchor score, in STRUCT (target mode). The process begins with an anchor and its associated targets, treating each as an anchor-targets simulation unit. The real dataset consists of anchors for which a stem-loop structure is detected in their base targets. For each anchor-targets simulation unit, nucleotides are randomly mutated at several positions in each target. In the example, the anchor has three targets, shown in the orange-shaded box. The green squares represent the stem locations, and the number of mismatches (v) in each target comes from the real data. For simulation, the corresponding v nucleotide positions in each target are randomly mutated (denoted as gray squares in the blue-shaded box), and a simulated *p*-value is computed. This process is repeated 1000 times for all anchor-targets units in the dataset. (B) Empirical cumulative distribution functions (ECDFs) of *p*-values (unadjusted) obtained from real data and 1000 simulated datasets. The ECDFs show unadjusted *p*-values obtained from the STRUCT output of a mosquito metatranscriptomics dataset (Batson et al.) alongside 1000 simulated datasets. Each data point corresponds to an anchor where a stem-loop is detected in the base compactor. The zoomed-in view highlights the enrichment of small *p*-values in the empirical dataset, indicating that STRUCT effectively identifies conserved stem-loops in RNA sequences.

**Suppl. Figure 5:**
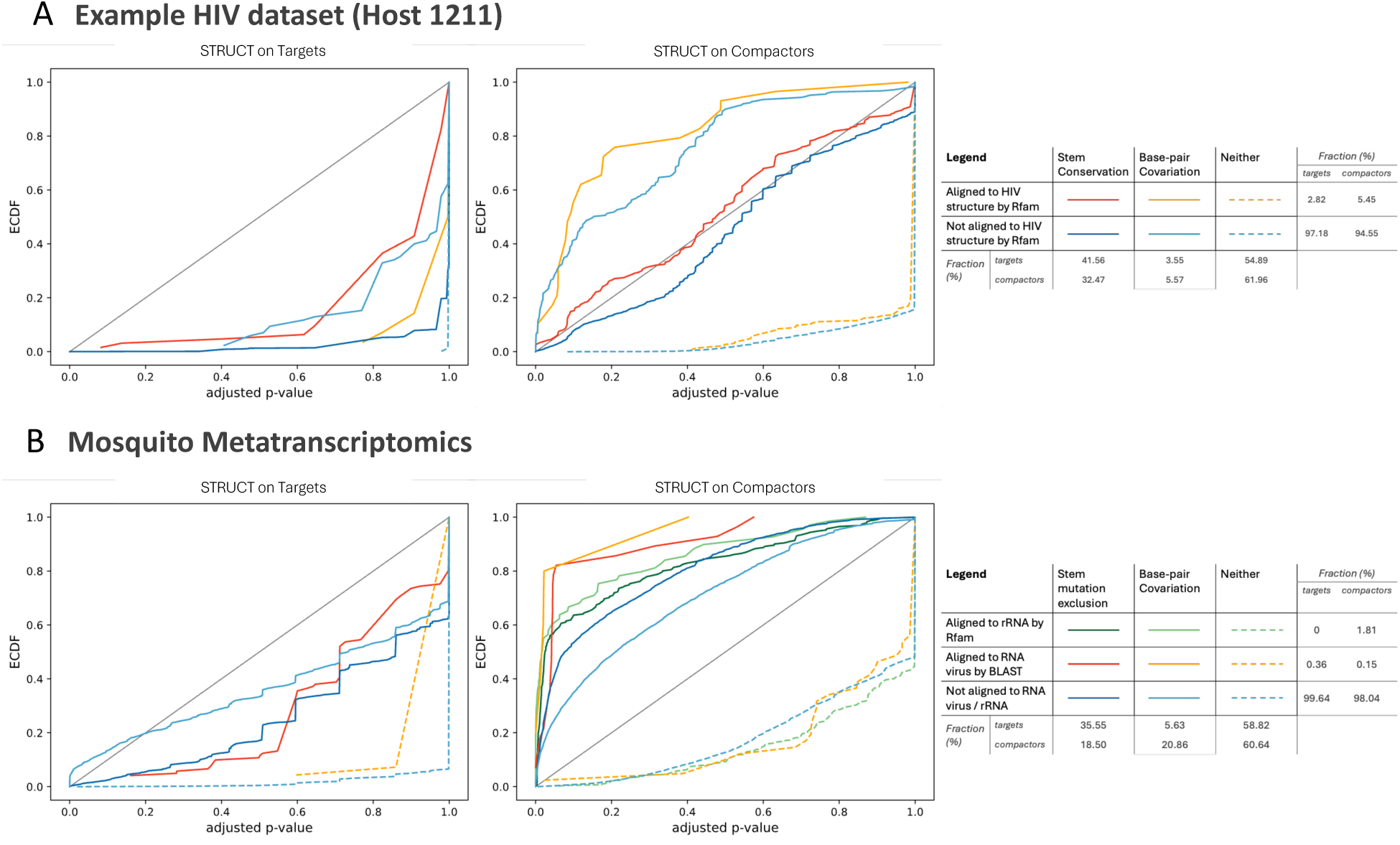
ECDF of BH-adjusted *p*-values in HIV and mosquito metatranscriptomics datasets. (A) and (B) show ECDF of adjusted *p*-values stratified across various RNA sequence categories obtained from different datasets using STRUCT on targets and compactors. For targets, each data point in the ECDF represents an anchor’s adjusted *p*-value. For compactors, each data point in the ECDF represents an anchor’s *p*-value for a way of split-and-concatenation of compactors. (A) For mosquito metatranscriptomics, all extendors (combined anchor and target sequences) and compactors undergo alignment with the Rfam database and a variety of viral genomes. Sequences that do not correspond with known rRNA sequences or match RNA virus genomes are categorized as “Unannotated RNA.” Sometimes different extendors (or compactors) linked to a single anchor get different annotations; for instance, if four out of five extendors associated with one anchor are identified as rRNA while one remains unannotated, we apply a majority rule approach for labeling the anchor, in this case as rRNA. The sequences are also classified based on the type of structural variation they exhibit, which is detailed in Figure 1C. The categories include BPC, SVE, and sequences with “Neither” (noting no conservation nor covariation in their stem regions). Similarly, the label for each anchor is determined by the majority annotation of its data points. (B) In the dataset of HIV, all extendors and compactors are aligned to the Rfam database. Sequences are classified by whether they align to an HIV structural element and by mechanisms of conserved structures. We employed the same majority rule to label anchors.

## References

[1] Rivas, E.: Evolutionary conservation of RNA sequence and structure 12(5), 1649 10.1002/wrna.1649. Accessed 2024-02-12

[2] Gundelfinger, E.D., Di Carlo, M., Zopf, D., Melli, M.: Structure and evolution of the 7sl RNA component of the signal recognition particle. 3(10), 2325–2332 10.1002/j.1460-2075.1984.tb02134.x. Accessed 2024-02-09

[3] Breaker, R.R.: Riboswitches and the RNA world 4(2), 003566–003566 10.1101/cshperspect.a003566. Accessed 2024-02-22

[4] Jackson, R.N., Golden, S.M., Van Erp, P.B.G., Carter, J., Westra, E.R., Brouns, S.J.J., Van Der Oost, J., Terwilliger, T.C., Read, R.J., Wiedenheft, B.: Crystal structure of the CRISPR RNA–guided surveillance complex from Escherichia coli 345(6203), 1473–1479 10.1126/science.1256328. Accessed 2024-06-24

[5] Peng, Y., Soper, T.J., Woodson, S.A.: RNase footprinting of protein binding sites on an mRNA target of small RNAs 905, 213–224 10.1007/978-1-61779-949-513

[6] Deigan, K.E., Li, T.W., Mathews, D.H., Weeks, K.M.: Accurate SHAPE-directed RNA structure determination 106(1), 97–102 10.1073/pnas.0806929106. Publisher: Proceedings of the National Academy of Sciences. Accessed 2024-06-18

[7] Spitale, R.C., Incarnato, D.: Probing the dynamic RNA structurome and its functions 24(3), 178–196 10.1038/s41576-022-00546-w. Publisher: Nature Publishing Group. Accessed 2024-10-03

[8] Zhang, J., Fei, Y., Sun, L., Zhang, Q.C.: Advances and opportunities in RNA structure experimental determination and computational modeling 19(10), 1193–1207 10.1038/s41592-022-01623-y. Accessed 2024-02-22

[9] Charon, J., Olendraite, I., Forgia, M., Chong, L.C., Hillary, L.S., Roux, S., Kupczok, A., Debat, H., Sakaguchi, S., Tahzima, R., Nakagawa, S., Babaian, A., Abroi, A., Bejerman, N., Ben Mansour, K., Brown, K., Butkovic, A., Cervera, A., Charriat, F., Chen, G., Chiba, Y., De Coninck, L., Demina, T., Dominguez-Huerta, G., Dubrulle, J., Gutierrez, S., Harvey, E., Jayaraj Mallika, F.R., Karapliafis, D., Lim, S.J., Kasibhatla, S.M., Mifsud, J.C.O., Nishimura, Y., Ortiz-Baez, A.S., Raco, M., Rivero, R., Sadiq, S., Saghaei, S., San, J.E., Shaikh, H.M., Sieradzki, E.T., Sullivan, M.B., Sun, Y., Wille, M., Wolf, Y.I., Zrelovs, N., Neri, U.: Consensus statement from the first RdRp summit: advancing RNA virus discovery at scale across communities 4 10.3389/fviro.2024.1371958. Publisher: Frontiers. Accessed 2024-10-03

[10] Rivas, E.: RNA structure prediction using positive and negative evolutionary information 16(10), 1008387 10.1371/journal.pcbi.1008387. Accessed 2024-02-22

[11] Rivas, E., Clements, J., Eddy, S.R.: A statistical test for conserved RNA structure shows lack of evidence for structure in lncRNAs 14(1), 45–48 10.1038/nmeth.4066. Number: 1 Publisher: Nature Publishing Group. Accessed 2024-01-22

[12] Jumper, J., Evans, R., Pritzel, A., Green, T., Figurnov, M., Ronneberger, O., Tunyasuvunakool, K., Bates, R., Žídek, A., Potapenko, A., Bridgland, A., Meyer, C., Kohl, S.A.A., Ballard, A.J., Cowie, A., Romera-Paredes, B., Nikolov, S., Jain, R., Adler, J., Back, T., Petersen, S., Reiman, D., Clancy, E., Zielinski, M., Steinegger, M., Pacholska, M., Berghammer, T., Bodenstein, S., Silver, D., Vinyals, O., Senior, A.W., Kavukcuoglu, K., Kohli, P., Hassabis, D.: Highly accurate protein structure prediction with AlphaFold 596(7873), 583–589 10.1038/s41586-021-03819-2. Publisher: Nature Publishing Group. Accessed 2024-06-11

[13] Noller, H.F., Kop, J., Wheaton, V., Brosius, J., Gutell, R.R., Kopylov, A.M., Dohme, F., Herr, W., Stahl, D.A., Gupta, R., Waese, C.R.: Secondary structure model for 23s ribosomal RNA 9(22), 6167–6189 10.1093/nar/9.22.6167

[14] Cheng, C.Y., Kladwang, W., Yesselman, J.D., Das, R.: RNA structure inference through chemical mapping after accidental or intentional mutations 114(37), 9876–9881 10.1073/pnas.1619897114. Publisher: Proceedings of the National Academy of Sciences. Accessed 2024-06-11

[15] Gao, W., Yang, A., Rivas, E.: Thirteen dubious ways to detect conserved structural RNAs 75(6), 471–492 10.1002/iub.2694. Accessed 2024-02-22

[16] Garcia-Vallvé, S., Romeu, A., Palau, J.: Horizontal gene transfer in bacterial and archaeal complete genomes 10(11), 1719–1725 10.1101/gr.130000

[17] Sanjuán, R., Nebot, M.R., Chirico, N., Mansky, L.M., Belshaw, R.: Viral mutation rates 84(19), 9733–9748 10.1128/jvi.00694-10. Publisher: American Society for Microbiology. Accessed 2024-10-03

[18] Stav, S., Atilho, R.M., Mirihana Arachchilage, G., Nguyen, G., Higgs, G., Breaker, R.R.: Genome-wide discovery of structured noncoding RNAs in bacteria 19(1), 66 10.1186/s12866-019-1433-7. Accessed 2024-02-09

[19] Burrill, C.P., Westesson, O., Schulte, M.B., Strings, V.R., Segal, M., Andino, R.: Global RNA structure analysis of poliovirus identifies a conserved RNA structure involved in viral replication and infectivity 87(21), 11670–11683 10.1128/JVI.01560-13. Accessed 2024-06-10

[20] Andrews, R.J., O’Leary, C.A., Tompkins, V.S., Peterson, J.M., Haniff, H., Williams, C., Disney, M.D., Moss, W.N.: A map of the SARS-CoV-2 RNA structurome 3(2), 043 10.1093/nargab/lqab043. Accessed 2024-06-10

[21] Lee, E., Bujalowski, P.J., Teramoto, T., Gottipati, K., Scott, S.D., Padmanabhan, R., Choi, K.H.: Structures of flavivirus RNA promoters suggest two binding modes with NS5 polymerase 12(1), 2530 10.1038/s41467-021-22846-1. Publisher: Nature Publishing Group. Accessed 2024-06-10

[22] Lauring, A.S., Andino, R.: Quasispecies theory and the behavior of RNA viruses 6(7), 1001005 10.1371/journal.ppat.1001005. Accessed 2024-02-23

[23] Antar, A.A.R., Jenike, K.M., Jang, S., Rigau, D.N., Reeves, D.B., Hoh, R., Krone, M.R., Keruly, J.C., Moore, R.D., Schiffer, J.T., Nonyane, B.A.S., Hecht, F.M., Deeks, S.G., Siliciano, J.D., Ho, Y.-C., Siliciano, R.F.: Longitudinal study reveals HIV-1–infected CD4+ t cell dynamics during long-term antiretroviral therapy 130(7), 3543–3559 10.1172/JCI135953. Accessed 2024-02-12

[24] Watts, J.M., Dang, K.K., Gorelick, R.J., Leonard, C.W., Bess Jr, J.W., Swanstrom, R., Burch, C.L., Weeks, K.M.: Architecture and secondary structure of an entire HIV-1 RNA genome 460(7256), 711–716 10.1038/nature08237. Accessed 2024-01-29

[25] Lin, Z., Akin, H., Rao, R., Hie, B., Zhu, Z., Lu, W., Smetanin, N., Verkuil, R., Kabeli, O., Shmueli, Y., Santos Costa, A., Fazel-Zarandi, M., Sercu, T., Candido, S., Rives, A.: Evolutionary-scale prediction of atomic-level protein structure with a language model 379(6637), 1123–1130 10.1126/science.ade2574. Publisher: American Association for the Advancement of Science. Accessed 2024-06-10

[26] Chaung, K., Baharav, T.Z., Henderson, G., Zheludev, I.N., Wang, P.L., Salzman, J.: SPLASH: A statistical, reference-free genomic algorithm unifies biological discovery 186(25), 5440–545626 10.1016/j.cell.2023.10.028

[27] Kokot, M., Dehghannasiri, R., Baharav, T., Salzman, J., Deorowicz, S.: Scalable and unsupervised discovery from raw sequencing reads using SPLASH2, 1–7 10.1038/s41587-024-02381-2. Publisher: Nature Publishing Group. Accessed 2024-09-26

[28] Vicens, Q., Kieft, J.S.: Thoughts on how to think (and talk) about RNA structure 119(17), 2112677119 10.1073/pnas.2112677119. Publisher: Proceedings of the National Academy of Sciences. Accessed 2024-04-24

[29] Henderson, G., Gudys, A., Baharav, T., Sundaramurthy, P., Kokot, M., Wang, P.L., Deorowicz, S., Carey, A.F., Salzman, J.: Ultra-efficient, unified discovery from microbial sequencing with SPLASH and precise statistical assembly. bioRxiv. Pages: 2024.01.18.576133 Section: New Results. 10.1101/2024.01.18.576133. 10.1101/2024.01.18.576133v1 Accessed 2024-01-26

[30] Lu, K., Heng, X., Summers, M.F.: Structural determinants and mechanism of HIV-1 genome packaging 410(4), 609–633 10.1016/j.jmb.2011.04.029. Accessed 2024-06-10

[31] Ye, L., Gribling-Burrer, A.-S., Bohn, P., Kibe, A., Börtlein, C., Ambi, U.B., Ahmad, S., Olguin-Nava, M., Smith, M., Caliskan, N., Von Kleist, M., Smyth, R.P.: Short– and long-range interactions in the HIV-1 5’ UTR regulate genome dimerization and packaging 29(4), 306–319 10.1038/s41594-022-00746-2. Accessed 2024-06-10

[32] Batson, J., Dudas, G., Haas-Stapleton, E., Kistler, A.L., Li, L.M., Logan, P., Ratnasiri, K., Retallack, H.: Single mosquito metatranscriptomics identifies vectors, emerging pathogens and reservoirs in one assay 10, 68353 10.7554/eLife.68353

[33] Biziaev, N.S., Egorova, T.V., Alkalaeva, E.Z.: Dynamics of eukaryotic mRNA structure during translation 56(3), 382–394 10.1134/S0026893322030037. Accessed 2024-09-25

[34] Passmore, L.A., Coller, J.: Roles of mRNA poly(a) tails in regulation of eukaryotic gene expression 23(2), 93–106 10.1038/s41580-021-00417-y

[35] Childs-Disney, J.L., Yang, X., Gibaut, Q.M.R., Tong, Y., Batey, R.T., Disney, M.D.: Targeting RNA structures with small molecules 21(10), 736–762 10.1038/s41573-022-00521-4. Accessed 2024-06-18

[36] Bruner, K.M., Murray, A.J., Pollack, R.A., Soliman, M.G., Laskey, S.B., Capoferri, A.A., Lai, J., Strain, M.C., Lada, S.M., Hoh, R., Ho, Y.-C., Richman, D.D., Deeks, S.G., Siliciano, J.D., Siliciano, R.F.: Defective proviruses rapidly accumulate during acute HIV-1 infection 22(9), 1043–1049 10.1038/nm.4156. Accessed 2024-07-07

[37] Kalvari, I., Nawrocki, E.P., Ontiveros-Palacios, N., Argasinska, J., Lamkiewicz, K., Marz, M., Griffiths-Jones, S., Toffano-Nioche, C., Gautheret, D., Weinberg, Z., Rivas, E., Eddy, S.R., Finn, R., Bateman, A., Petrov, A.I.: Rfam 14: expanded coverage of metagenomic, viral and microRNA families 49, 192–200 10.1093/nar/gkaa1047. Accessed 2024-04-16

[38] Camacho, C., Coulouris, G., Avagyan, V., Ma, N., Papadopoulos, J., Bealer, K., Madden, T.L.: BLAST+: architecture and applications 10(1), 1–9 10.1186/1471-2105-10-421. Number: 1 Publisher: BioMed Central. Accessed 2024-04-16

[39] Nawrocki, E.P., Eddy, S.R.: Infernal 1.1: 100-fold faster RNA homology searches 29(22), 2933–2935 10.1093/bioinformatics/btt509. Accessed 2024-04-16

[40] Kerpedjiev, P., Hammer, S., Hofacker, I.L.: Forna (force-directed RNA): Simple and effective online RNA secondary structure diagrams 31(20), 3377–3379 10.1093/bioinformatics/btv372. Accessed 2024-05-31

