## Supplemental Methods for "STRUCT: a statistical approach to identify RNA secondary structures from raw sequencing data, bypassing multiple sequence alignment"

2024

### 1 Closed-form target p-value for base-pair covariation (BPC) test

In the BPC test, we define the conditional likelihood of having at least as many compensatory mutation pairs $C$ as the observed value $c$, given the predetermined parameters target length $k$, stem length $L$, and number of mismatches $v$, as the first target p-value, denoted as $p^{(1)}_{\text{target}}$,

$$\begin{aligned} p^{(1)}_{\text{target}} &= \mathbb{P}(C \geq c \mid L, v, k) \\ &= \sum_{h=2c}^{\min(v,2L)} \mathbb{P}(C \geq c \mid L, h)\, \mathbb{P}(H = h \mid L, v, k). \end{aligned} \tag{1}$$

where the last line is allowed by the law of total probability. $H$ denotes the variable representing the number of mismatches observed in stems, and $h$ denotes the possible values that $H$ can take on.

In the preceding equation, the second term can be modeled with a hypergeometric distribution,

$$\mathbb{P}(H = h \mid L, v, k) = \frac{\binom{2L}{h}\binom{k-2L}{v-h}}{\binom{k}{v}}. \tag{2}$$

The first term can be decomposed into the summation of discrete cases of $g$ pairs of compensatory mutations in a target.

$$\mathbb{P}(C \geq c \mid L, h) = \sum_{g=c}^{\lfloor h/2 \rfloor} \mathbb{P}(g \mid L, h) \tag{3}$$

Moreover, denote $l$ as the number of mismatches in the left stem. We can write $\mathbb{P}(g \mid L, h)$ in a closed form:

$$\begin{aligned} &\mathbb{P}(g \mid L, h) \\ &= \sum_{l=g}^{\min(L,h)} \frac{\binom{L}{l'} 3^{l'} \binom{l'}{g} \sum_{m=0}^{l'-g} \binom{l'-g}{m} 2^m \binom{L-l'}{h-l'-g-m} 3^{h-l'-g-m}}{\binom{2L}{h} 3^h} \end{aligned} \tag{4}$$

where $l' = \min(h-l, l)$.

### 2 String notation

STRUCT outputs a string notation to visualize the composition of each tested sequence, whether it's a target or a sub-sequence from a compactor, compared to its base target or the sub-sequence in the base compactor, in which a stem-loop is detected. For example, consider the following base target and a varied target:

```
Base target:         TACCACTTTAA ATGGC GAACA GCCAT A
Target 1:            TACCGCTTTAA TGGGC GTACA ACCCA A
Structure notation:  ----g------{TG---(-t---)a--CA}-
```

In the structure notation, each dash or letter corresponds to a nucleotide in the tested sequence. The braces {} enclose the predicted stem-loop structure, with the parentheses () containing the predicted loop. Sequences flanked by { and ( or by } and ) are stems. Within the stems, capital letters denote predicted compensatory mutations, while lowercase letters indicate predicted non-compensatory mutations. In the whole sequence, nucleotides matching the base sequence are shown as dashes, while those mutated relative to the base sequence appear in lowercase.

Under compactor mode, if the stem-loop structure is only detected in one of the two recombined sub-sequences, only one notation is output.

### 3 *forna* RNA structure rendering plots

*forna* package provides a web interface for visualizing RNA secondary structures. We provide sequences and the predicted secondary structures in dot-bracket notation as input, as well as a custom color scheme denoting compensatory mismatches and mismatches. For example, a forna input looks like:

```
> base_target
TACCACTTTAAATGGCGAACAGCCATA
...........(((((.....))))).
> target_1
TACCGCTTTAATGGGCGTACAACCCAA
...........(((((.....))))).
```

The custom color scheme is

```
>base-target
1-27:lightblue 12-16,22-26:lightgreen
>target-1
1-27:lightblue 12-16,22-26:lightgreen 5,18,22:grey 12,13,25,26:
    lightsalmon
```
